# Insulin/IGF signaling and TOR promote vitellogenesis via inducing juvenile hormone biosynthesis

**DOI:** 10.1101/2020.04.06.028639

**Authors:** Shiming Zhu, Fangfang Liu, Huanchao Zeng, Na Li, Chonghua Ren, Yunlin Su, Shutang Zhou, Guirong Wang, Subba Reddy Palli, Jian Wang, Yiru Qin, Sheng Li

## Abstract

Vitellogenesis, including vitellogenin (Vg) production in the fat body and Vg uptake by maturing oocytes, is of great importance for the successful reproduction of adult females. The endocrinal and nutritional regulation of vitellogenesis differs distinctly in insects. Here, the complex crosstalk between juvenile hormone (JH) and the two nutrient sensors, insulin/IGF signaling (IIS) and target of rapamycin (TOR), was investigated to elucidate the molecular mechanisms regulating vitellogenesis in the American cockroach, *Periplaneta americana.* Our data showed that a block of JH biosynthesis or JH action arrested vitellogenesis, partially by inhibiting the expression of *doublesex (Dsx),* a key transcription factor gene involved in the sex determination cascade. Depletion of IIS and TOR blocked both JH biosynthesis and vitellogenesis. Importantly, the JH analog methoprene, but not bovine insulin (to restore IIS) and amino acids (to restore TOR activity), restored vitellogenesis in the neck-ligated (both nutrient- and JH-deficient) cockroaches. Combining classic physiology with modern molecular techniques, we have demonstrated that JH signaling alone is able to induce vitellogenesis and thus ovarian maturation. By contrast, IIS and TOR do not induce vitellogenesis independent of JH, the nutrient sensors promote vitellogenesis in an indirect manner via activating JH biosynthesis.

## INTRODUCTION

Vitellogenesis, an imperative event of insect reproduction, involves vitellogenin (Vg) production in the fat body, the release of Vg into the hemolymph, and Vg uptake by and its modification in maturing oocytes. During the process of Vg uptake, extensive intercellular spaces are formed among follicle cells (follicular patency), allowing the internalization of hemolymph Vg into maturing oocytes by receptor-mediated endocytosis (Raikhel, 2005; Roy et al., 2018). However, the connection between Vg production and the formation of follicular patency has not been determined yet.

Juvenile hormone (JH), a structurally unique sesquiterpenoid hormone found only in arthropods, plays prominent roles in the regulation of reproduction, metamorphosis, and other biological processes (Li et al. 2019; Jindra et al., 2013; Roy et al., 2018). JH is primarily biosynthesized in the corpora allata (CA) of insects through 13 discrete enzymatic steps (Bellés et al., 2005). The later two steps of JH biosynthesis are catalyzed by two crucial regulatory enzymes, juvenile hormone acid methyltransferase (Jhamt) and methyl farnesoate epoxidase (Cyp15a1) (Defelipe et al., 2011; Shinoda and Itoyama, 2003). JH signal transduction involves both genomic and nongenomic actions. In the genomic action, JH binds its intracellular receptor, methoprene-tolerant (Met), to induce the expression of a JH primary-response gene *Krüppel-homolog 1 (Kr-h1)*, which plays a central role in JH signaling (Jindra et al., 2013; Li et al., 2019). Meanwhile, JH signaling may be transduced via receptor tyrosine kinase (RTK) or G protein-coupled receptor (GPCR), putative plasma membrane receptors, to act in a nongenomic manner (Davey, 2000; Liu et al., 2015; Ojani et al., 2016; Bai and Palli, 2016). JH is one of the main gonadotrophic hormones that promote vitellogenesis in insects, including Vg production in the fat body via the genomic action as well as the development of follicular patency and thus Vg uptake by the maturing oocytes via the non-genomic action (Raikhel et al., 2005; Jing et al., 2018; Roy et al., 2018).

*Doublesex* (*Dsx*), a transcription factor in the last step of the sex determination cascade, participates in reproductive regulation in female insects (Verhulst and Van de Zande, 2015). Dsx was reported to induce *Vg* expression in the fruit fly *Drosophila melanogaster* (Burtis et al., 1991), the wild silkmoth *Antheraea assama* (Shukla and Nagaraju, 2010), and the red flour beetle *Tribolium castaneum* (Shukla and Palli, 2012). Moreover, a causal link between Dsx and JH signaling was revealed in the stag beetle *Cyclommatus metallifer* (Gotoh et al., 2014). Therefore, *Dsx* is probably a transcription factor involved in the JH regulation of vitellogenesis.

Insulin/IGF signaling (IIS) and target of rapamycin (TOR) are highly evolutionarily conserved nutrient sensors with vital key regulatory roles in development and diseases (Li et al., 2019b; Roy et al., 2018). Insulin/IGF binds the membrane insulin receptor (InR) and activates IIS by a series of protein phosphorylation events, leading to the activation of protein kinase B (Akt) (Nässel and Broeck, 2016; Li et al., 2019b). TOR receives IIS and amino acid signals, phosphorylates effector proteins such as 4EBP and S6K to stimulate protein synthesis, ribosome biogenesis, and cell growth (Li et al., 2019b; Smykal and Raikhel, 2015). It is well known that a block of IIS or TOR decreases JH biosynthesis in the CA and *Vg* expression in the fat body in a number of insect species. However, the interplay between the nutrient sensors and JH signaling in the regulation of vitellogenesis varies dramatially in insects (Abrisqueta et al., 2014; Corona et al., 2007; Guo et al., 2014; Li et al., 2018; Maestro et al., 2009; Parthasarathy and Palli, 2011; Parthasarathy et al., 2010; Perez-Hedo et al., 2013; Sheng et al., 2011; Song et al., 2014). Nevertheless, more solid evidence is indispensable to demonstrate whether IIS and TOR induce vitellogenesis independent of JH or act indirectly through regulating JH biosynthesis.

The American cockroach *Periplaneta americana* is an invasive urban pest in both warm and humid regions, partially due to its high fecundity. Our previous work showed that JH signaling, IIS, and TOR, and Dsx affect ovarian maturation in American cockroach (Li et al., 2018). However, how these signals interact to promote vitellogenesis remains unknown. In this study, we have determnined that IIS and TOR promote vitellogenesis in an indirect manner by activating JH biosynthesis. Combining classic physiology with morden molecular techniques, our studies broaden the understanding of endocrinal and nutritional regulation of female reproduction in insects.

## RESULTS

### Vg production is required for development of functional follicular patency

Under our rearing conditions, the first reproductive cycle of adult female American cockroaches was approximately 8 days, and the primary oocytes start to accumulate yolk proteins (mainly Vg) at approximately day 4 post-adult emergence (PAE) (Li et al., 2018). Protein mass spectroscopy analysis revealed that the ovarian proteins in the 100 kDa major band are mainly the two Vg proteins, Vg1 and Vg2 (Fig. S1A, B). Since this band was most abundant and no antibody against Vg was available at present, it was used to quantify the ovarian Vg content in this study. The Vg protein levels were low during the first 3 days PAE, reached the maximum levels on day 5 PAE, and remained stable until oviposition (day 8 PAE) (Fig. 1A, A’). The results of quantitative real-time PCR (qPCR) revealed that the *Vg*1 transcription in the fat body began on day 3 PAE, peaked at the middle cycle, and steadily decreased thereafter (Fig. 1A’), showing a similar but slightly advanced developmental pattern in comparison with Vg protein levels in the ovary.

**Figure 1.**
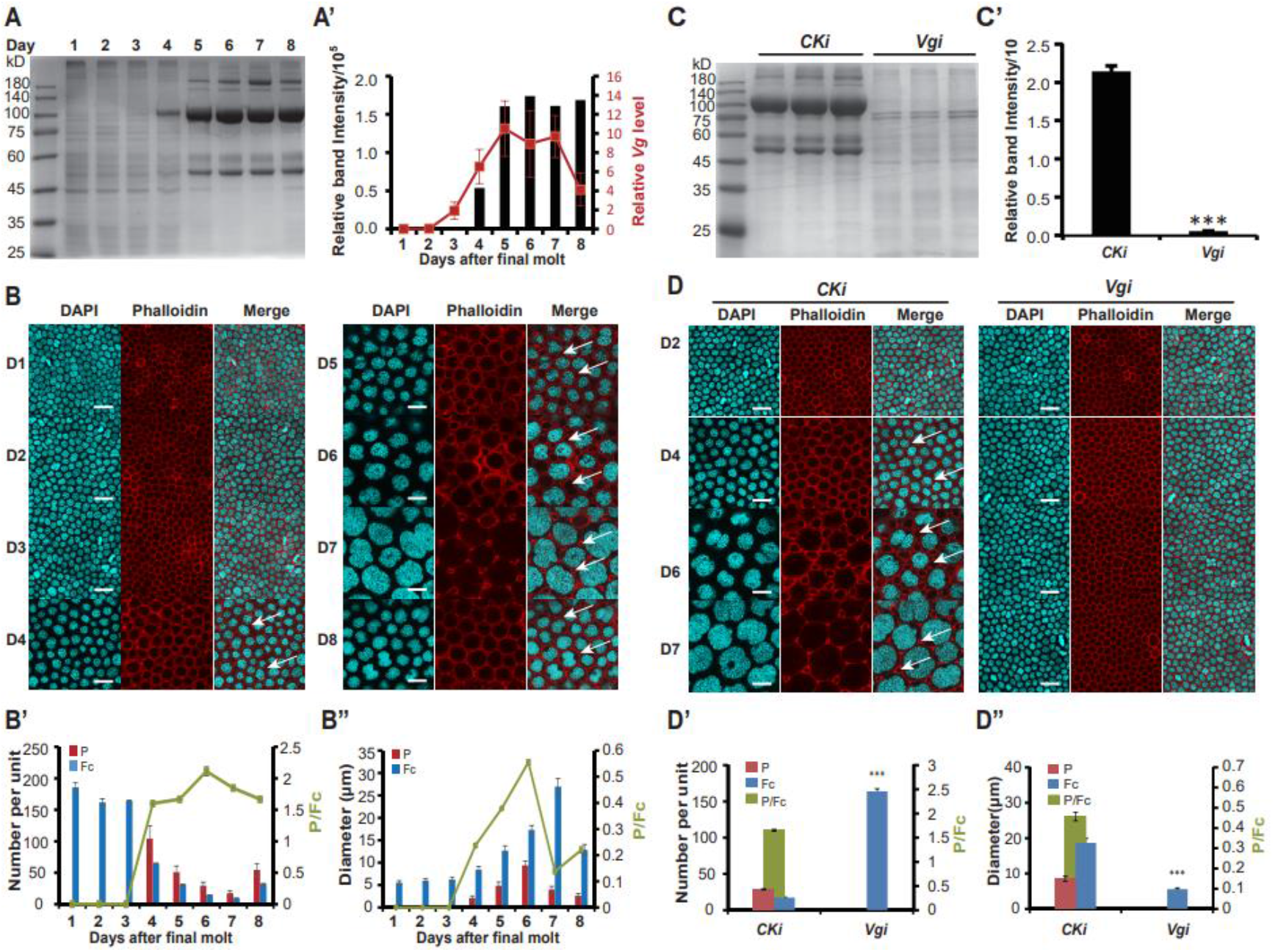
Vg production is required for development of functional follicular patency. **(A, A’)** (A) Temporal changes of total protein in the ovary. The total protein was separated by SDS-PAGE. Mass spectrometry analysis showed that the 100 kDa proteins were Vg1 and Vg2 (Fig. S1); (A’) quantification of the 100 kDa band intensity in (A) and *vitellogenin (Vg)* mRNA level in the fat body from day 1 to day 8 PAE. n=3. **(B-B”)** (B) Morphological change of follicular epithelium from day 1 to day 8 PAE; (B’) Quantification of follicle cells (Fc) and patency (P) number in a unit and the number index (P/Fc); (B”) Quantification of the diameter of Fc nuclei and P and the diameter index (P/Fc). Blue, follicle cell nuclei; red, F-actin. White arrows indicate follicular patency. Scale bar: 20 μm. n=3. **(C, C’)** (C) Effect of *Vg* RNAi on Vg protein levels in the ovary; (C’) The quantification of 100 kDa band intensity showing a decrease in Vg protein levels after *Vg* RNAi. n=3. **(D-D”)** (D) Effect of *Vg* RNAi on the follicular epithelium development; (D’) Quantification of Fc and P number in a unit and the number index (P/Fc); (D”) Quantification of the diameter of Fc nuclei and P and the diameter index (P/Fc). Blue, follicle cell nuclei; red, F-actin. n=3. White arrows indicate patency. Scale bar: 20 μm. \*\*\**P*<0.001.

The developmental profiles of the sizes of the follicle cells and follicular patency in the maturing oocytes were investigated to monitor ovarian maturation. The size of follicle cells remained constant during the first 3 days PAE, increased significantly from day 3 to day 7 PAE, and reduced slightly at day 8 PAE. Follicular patency appeared at day 4 PAE, indicating that the oocyte started to take up Vg. Although the number of follicular patency decreased in a unit, its diameter linearly increased from day 3 to day 6 PAE, and decreased thereafter (Fig. 1B-B”). Therefore, the number index (patency number/ follicle cells number) profile remained constantly high after day 4 PAE, while the diameter index (patency diameter/ follicle cell nuclei diameter) profile mirrored Vg production (Fig. 1B-B”). This quantitative technique serves as an effective method to evaluate the maturation of the ovary.

Because the *Vg*1 DNA fragment used to design *Vg*1 dsRNA showed a relatively high degree of similarity to the related *Vg*2 fragment, RNA interference (RNAi) using *Vg*1 dsRNA significantly reduced the expression of both *Vg1* and *Vg*2 (Fig. S1C, D). *Vg* RNAi reduced not only *Vg* mRNA expression in the fat body (Fig. S1C) and Vg protein accumulation in the ovary (Fig. 1C, C’), but also the development of follicle cells and follicular patency (Fig. 1D-D”; S1E, F). These results demonstrated that Vg production is required for the development of follicular patency and ovarian maturation.

### JH signaling regulates vitellogenesis partially by inducing *Dsx* expression

To reveal the possible relationship of Dsx, Met, and Kr-h1, we first analyzed their gene expression profiles in the fat body during the first reproductive cycle. The developmental profiles of *Dsx, Met,* and *Kr-h1* were analogous to that of *Vg1,* while *Dsx* showed a slightly precocious expression profile compared to *Met* and *Kr-h1* (Fig. 2A, B). A 95% decrease in *Dsx* mRNA levels were detected in the fat body of females injected with *Dsx* dsRNA, and *Vg* mRNA levels were reduced by 71.8% by *Dsx* RNAi (Fig. 2C). By contrast, *Vg* RNAi had no effects on *Dsx* expression (Fig. 2C). The results together suggest the dependence of *Vg* expression on *Dsx* transcription.

**Figure 2.**
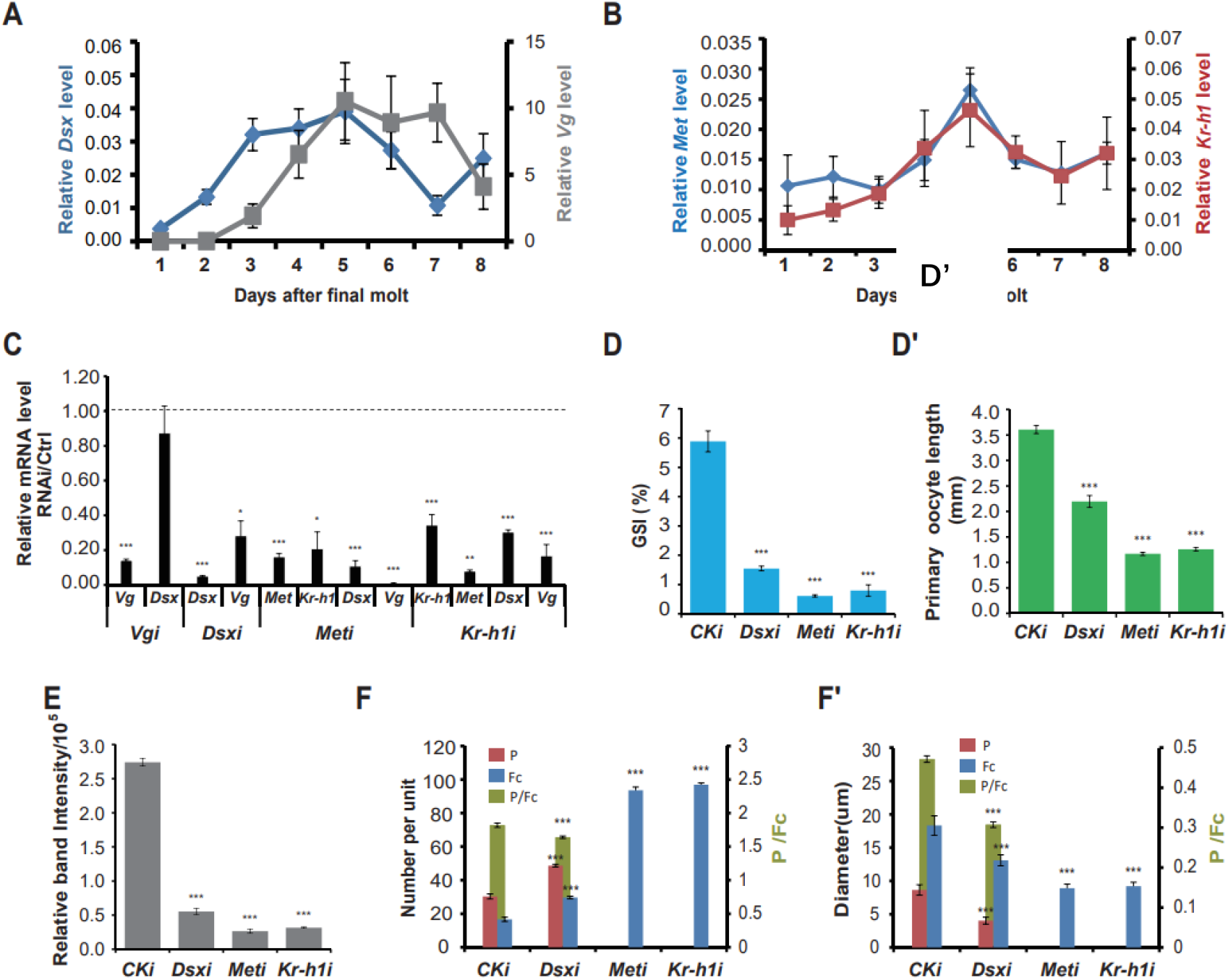
JH signaling regulates vitellogenesis partially by inducing *Dsx* expression. **(A, B)** Expression profiles of *Dsx*, *Vg* (A) and *Met*, *Kr-h1* (B) from day 1 to day 8 PAE. n=4. **(C-F’)** Effects of *Vg* RNAi, *Dsx* RNAi, *Met* RNAi, or *Kr-h1* RNAi on the expression of *Vg*, *Dsx*, *Met,* and *Kr-h1* (C), the gonadosomatic index (GSI) (D), the primary oocyte length (D’), the 100 kDa band intensity (E), number of Fc and P and the number index (P/Fc) (F), diameter of Fc nuclei and P and the diameter index (P/Fc) (F’). n=4. \**P*<0.05, \*\**P*<0.01, \*\*\**P*<0.001, compared to the negative control (CK RNAi).

Upon dsRNA treatments, *Met* and *Kr-h1* mRNA levels in the fat body were significantly reduced (Fig. 2C). Interestingly, *Met* RNAi decreased the *Kr-h1* mRNA level, while *Kr-h1* RNAi also decreased the *Met* mRNA level (Fig. 2C), indicating that *Kr-h1* is downstream of Met and Kr-h1 may regulate *Met* expression through a positive feedback loop. Knockdown of *Met* and *Kr-h1* led to a significant reduction in *Dsx* and *Vg* mRNA levels in the fat body (Fig. 2C); thus, the expression of *Dsx* and *Vg* depends on JH signaling.

Importantly, RNAi depletion of *Dsx*, *Met*, or *Kr-h1* blocked the maturation of primary oocytes and the follicular epithelium (Fig. S2A-C) and significantly reduced the gonadosomatic index (GSI) (Fig. 2D), primary oocyte length (Fig. 2D’), accumulation of Vg in the ovary (Fig. 2E), the diameter of follicle cells and follicular patency (Fig. S2D), the number index (Fig. 2F), and the diameter index (Fig. 2F’). The above characteristic altered in the maturing oocytes together were used to monitor vitellogenesis and ovarian maturation in the following. It is necessary to note that, in contrast with the *Met-* or *Kr-h1-*depleted cockroaches in which neither ovarian growth nor follicular patency was observed, a certain amount of Vg still accumulated in the primary oocytes and size-reduced follicular patency still occured in the *Dsx-*depleted cockroaches, implying that Dsx is not the single transcription factor downstream of JH signaling in the regulation of vitellogenesis. These experimental data revealed that JH signaling regulates vitellogenesis partially by inducing *Dsx* expression.

### JH biosynthesis is required for vitellogenesis

The mRNA abundance of *Jhamt* and *Cyp15a1* in the CA during the first reproductive cycle was determined by qPCR. The expression of the two JH biosynthetic enzyme genes increased from day 3 PAE, peaked on day 4-5 PAE, and decreased thereafter (Fig. 3A), showing similar developmental patterns to JH signaling and vitellogenesis. The effects of JH biosynthesis on vitellogenesis were examined by RNAi knockdown of *Jhamt* or *Cyp15a1.* RNAi depletion of *Jhamt* or *Cyp15a1* in the CA significantly suppressed the mRNA expression of *Met*, *Kr-h1, Dsx,* and *Vg* in the fat body (Fig. 3B). Moreover, the depletion of either of *Jhamt* or *Cyp15a1* blocked vitellogenesis and ovarian maturation (Fig. 3C-E’; S3). These data show that JH biosynthesis is required for JH signaling, which then promotes vitellogenesis.

**Figure 3.**
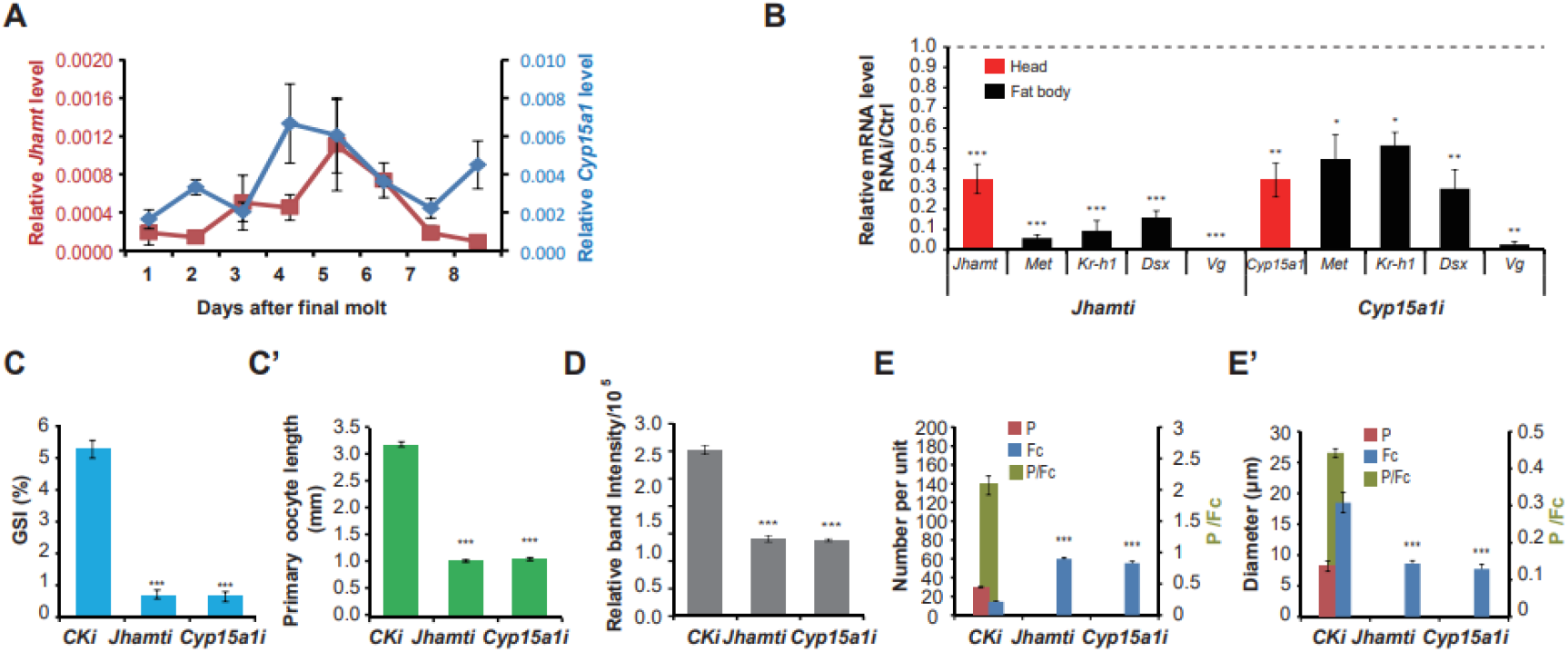
JH biosynthesis is required for vitellogenesis. **(A)** Expression profiles of *Jhamt* and *Cyp15a1* genes from day 1 to day 8 PAE. n=4. **(B-E’)** Effects of *Jhamt* RNAi and *Cyp15a1* RNAi on the expression of *Jhamt* and *Cyp15a1, Vg, Dsx, Met,* and *Kr-h1* (B) (the mRNA of *Jhamt* and *Cyp15a1* was extracted from heads, while the mRNA of *Met*, *Kr-h1, Dsx,* and *Vg* was extracted from fat body), the gonadosomatic index (GSI) (C), the primary oocyte length (C’), the 100 kDa band intensity (D), number of Fc and P and the number index (P/Fc) (E), diameter of Fc nuclei and P and the diameter index (P/Fc) (E’). n=4. \**P*<0.05, \*\**P*<0.01, \*\*\**P*<0.001, compared to the negative control (CK RNAi).

Next, we investigated the possible roles of the nutrient sensors (IIS and TOR) in the promotion of vitellogenesis and the interplay between the nutrient sensors and JH.

### IIS and TOR are required for JH biosynthesis and vitellogenesis

We thus measured the developmental profiles of IIS and TOR in the fat body during the first reproductive cycle. The total Akt protein level in the first reproductive cycle remained quite constant, while the level of phosphorylated Akt (P-Akt, refecting IIS) increased gradually until the middle cycle and slightly decreased thereafter (Fig. 4A). The effects of IIS on vitellogenesis were determined by reducing IIS with *InR* RNAi or LY294002 (a PI3K inhibitor) treatment. Both approaches significantly decreased P-Akt levels and thus IIS (Fig. 4A, A’). RNAi Knockdown of *InR* caused a significant reduction in the expression of *Jhamt*, *Cyp15a1*, *Met*, *Kr-h1*, *Dsx*, and *Vg*, which was comparable to the effects of LY294002 treatment (Fig. 4B), indicating that IIS regulates JH biosynthesis and the expression of *Dsx* and *Vg*. While vitellogenesis and ovarian maturation in the control animals progressed, they were blocked by *InR* dsRNA or LY294002 treatment (Fig. 4C-F”), indicating that IIS is required for both JH biosynthesis and vitellogenesis.

**Figure 4.**
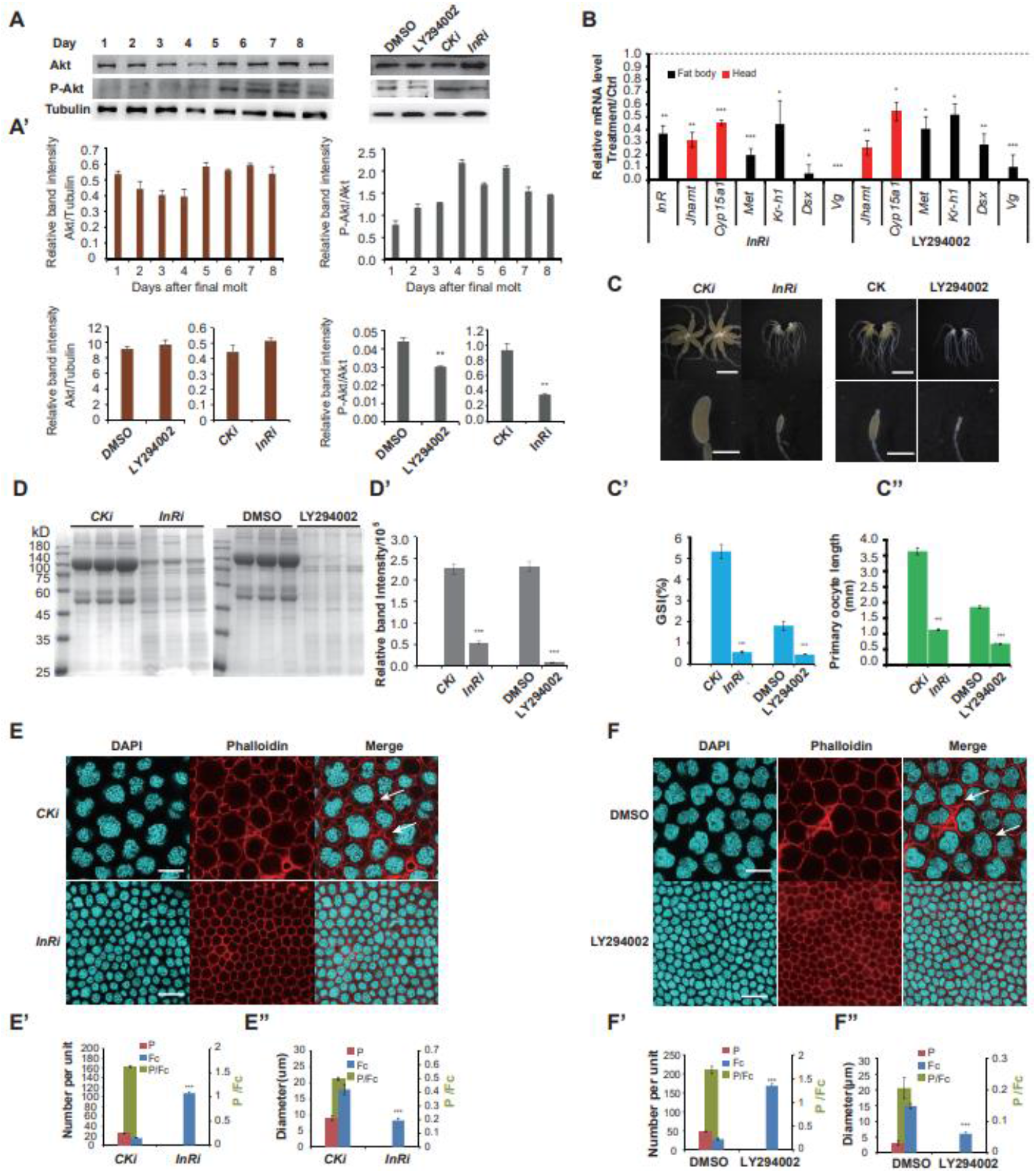
IIS is required for JH biosynthesis and vitellogenesis. **(A, A’)** Protein levels of Akt and phosphorylated Akt (P-Akt, reflecting IIS) in the fat body. (A) Temporal changes of the protein levels of Akt and P-Akt from day 1 to day 8 PAE (left panel) and the effects of IIS depletion (treatments with *InR* dsRNA and LY294002) on the protein levels of Akt and P-Akt (right panel). (A’) Quantification of band intensity showing the decrease of P-Akt protein levels after *InR* RNAi and LY294002 treatment. n=3. **(B-F”)** Effects of *InR* dsRNA and LY294002 on the expression of *Jhamt* and *Cyp15a1, Met, Kr-h1, Dsx,* and *Vg* (the mRNA of *Jhamt* and *Cyp15a1* was extracted from heads, while the mRNA of *Met*, *Kr-h1, Dsx,* and *Vg* was extracted from fat body) (B), the morphology of ovary, the gonadosomatic index (GSI), and the primary oocyte length (C-C’), the 100 kDa band intensity (D, D’), number of Fc and P and the number index (P/Fc) (E-E”), diameter of Fc nuclei and P and the diameter index (P/Fc) (F-F”). Scale bar: 20 μm. n=4. \**P*<0.05, \*\**P*<0.01, \*\*\**P*<0.001, compared to the negative control (CK RNAi or DMSO).

Similar to the level of P-Akt, the level of phosphorylated S6K (P-S6K, reflecting TOR) gradually increased until day 4 PAE and then slightly decreased in the first reproductive cycle (Fig. 5A, A’). Slimfast (Slif) is a membrane transporter of amino acids involved in TOR activation (Colombani et al., 2003). We injected cockroaches with *Slif* dsRNA, *TOR* dsRNA, or rapamycin (a TOR inhibitor) to examine the effects of TOR on vitellogenesis. Both rapamycin treatment and the RNAi knockdown of *Slif* or *TOR* decreased P-S6K levels and thus TOR activity (Fig. 5A, B). Compared to the control, the mRNA levels of *Jhamt*, *Cyp15a1*, *Met*, *Kr-h1*, *Dsx*, and *Vg* were significantly decreased after TOR activity was blocked (Fig. 5B). Furthermore, the block of TOR activity arrested vitellogenesis and ovarian maturation (Fig. 5C-F’; S4 A-F). These data indicate that, the same as IIS, TOR activity is a prerequisite for both JH biosynthesis and vitellogenesis.

**Figure 5.**
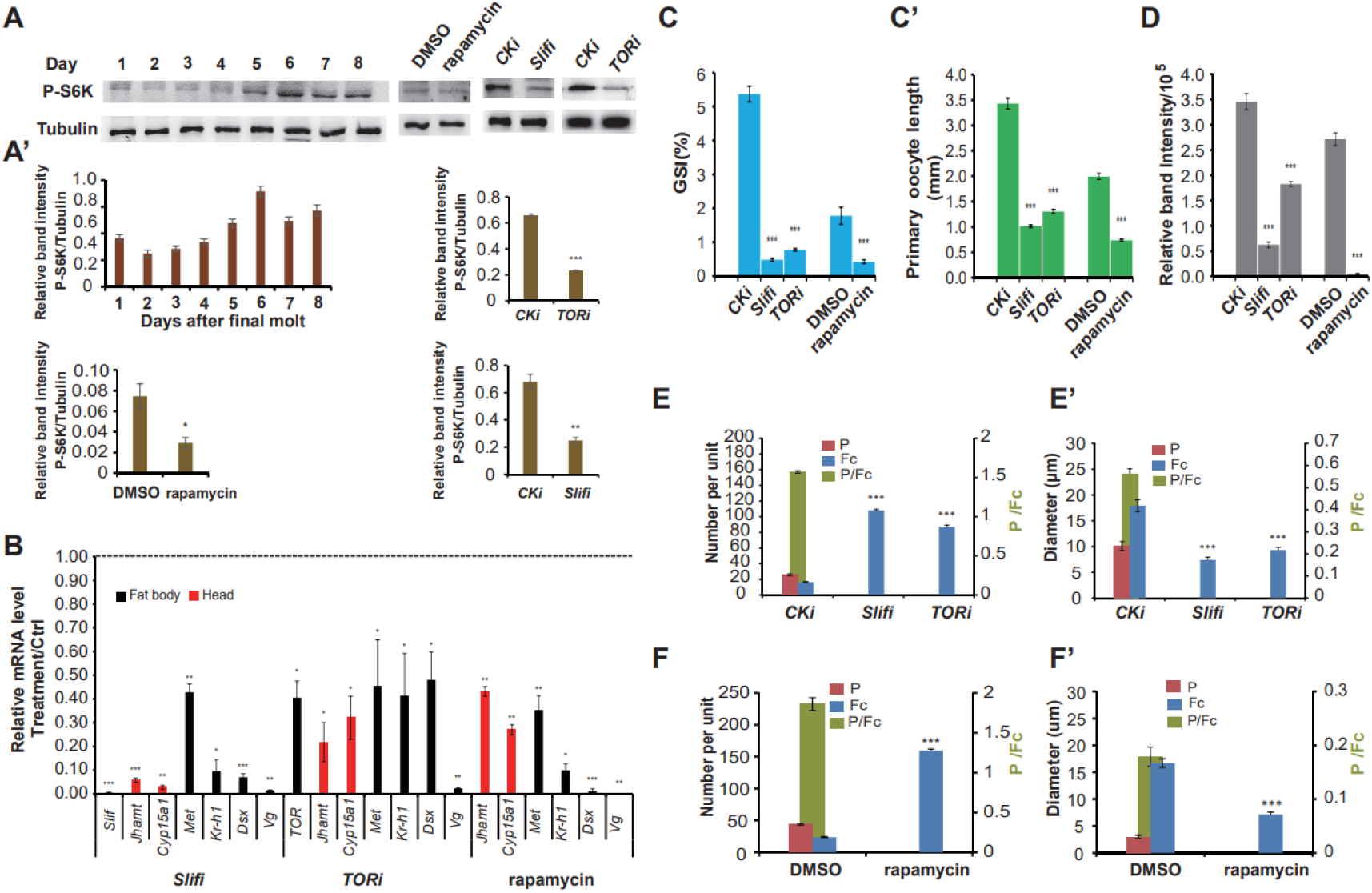
TOR is required for JH biosynthesis and vitellogenesis. **(A, A’)** Protein levels of phosphorylated S6K (P-S6K, reflecting TOR) in the fat body. (A) Temporal changes of the protein levels of p-S6K from day 1 to day 8 PAE (left panel) and the effects of TOR depletion (treatments with *Slif* dsRNA, *TOR* dsRNA, and rapamycin) on the P-S6K protein levels of (right panel). (A’) Quantification of band intensity showing the decrease of P-S6K protein levels after the treatments.. **(B-F”)** Effects of *InR* dsRNA and LY294002 on the expression of *Jhamt* and *Cyp15a1*, *Met*, *Kr-h1*, *Dsx*, and *Vg* (the mRNA of *Jhamt* and *Cyp15a1* was extracted from heads, while the mRNA of *Met*, *Kr-h1, Dsx,* and *Vg* was extracted from fat body) (B), the gonadosomatic index (GSI) and the primary oocyte length (C-C’), the 100 kDa band intensity (D), number of P and Fc and the number index (P/Fc) (E, F), diameter of Fc nuclei and P and the diameter index (P/Fc) (E’, F’). n=4. \**P*<0.05, \*\**P*<0.01, \*\*\**P*<0.001, compared to the negative control (CK RNAi or DMSO).

### JH signaling promotes vitellogenesis independent of IIS and TOR

We then examined the effects of starvation (nutrient-deficient) and neck ligation (both nutrient- and JH-deficient) on ovarian maturation during the first reproductive cycle. Interestingly, the ovaries still developed to a certain extent after starvation, while ovarian development was completely arrested after neck ligation, implying JH alone might promote vitellogenesis and ovarian maturation (Fig. 6A-A”). Based on these data, we chose the neck ligation approach for the following rescue experiments by injections of a JH analog methoprene, bovine insulin, and an amino acid mixture to restore JH signaling, IIS, and TOR activity, respectively.

**Figure 6.**
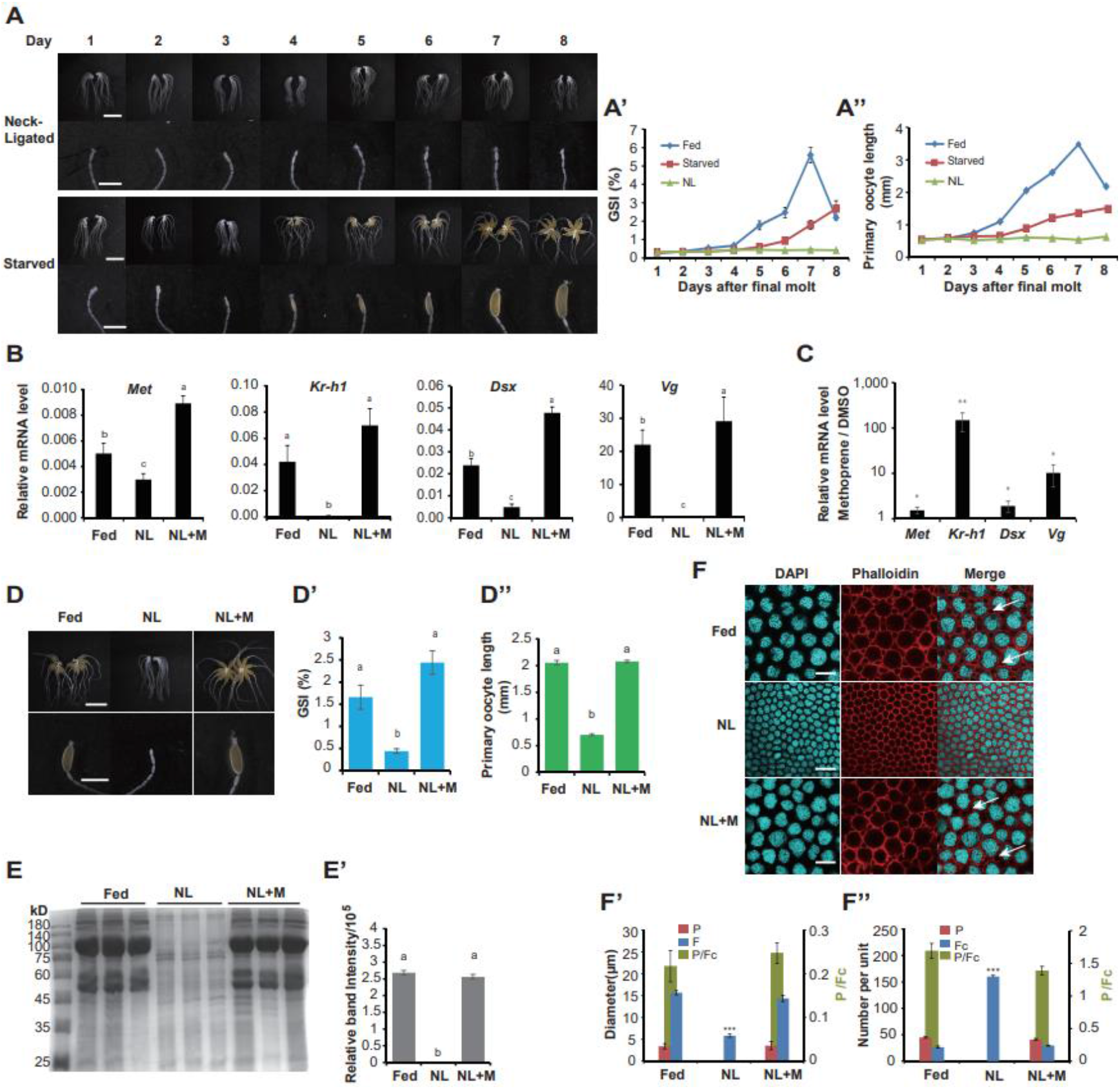
JH signaling promotes vitellogenesis independent of IIS and TOR. **(A-A”)** Themporal changes of ovary from day 1 to day 8 PAE after neck ligation (NL) and starvation. Morphological changes (A), the quantification of GSI (A’), the quantification of primary oocyte length (A”). **(B)** Effects of neck ligation (NL) and subsequent rescue with methoprene (a JH analog) (NL+M) on the expression of *Vg*, *Dsx*, *Met*, and *Kr-h1* in the fat body. **(C)** Effects of methoprene treatment on the expression of *Vg*, *Dsx*, *Met*, and *Kr-h1* on the *in vitro* cultured fat body. **(D-F”)** Effects of NL and NL+M on morphology of ovary, the gonadosomatic index (GSI) and the primary oocyte length (D-D”), the 100 kDa band intensity (E, E’), number of Fc and P and the number index (P/Fc), diameter of Fc nuclei and P and the diameter index (P/Fc) (F-F”). Blue, follicle cell nuclei; red, F-actin. n=3. White arrows indicate patency. Scale bar: 20 μm. \*\*\**P*<0.001. The bars labeled with different letters indicate significant difference at *P*< 0.05.

Neck ligation caused a significant reduction in *Met*, *Kr-h1, Dsx,* and *Vg* mRNA levels in the fat body compared to their levels in the feeding group. Importantly, methoprene treatment restored the expression of these genes to the levels even higher than those in the feeding group (Fig. 6B). To further validate these results, an *in vitro* culture experiment was carried out. Fat body from newly emerged females *(Vg* expression was not detected) were *in vivo* incubated for 6 h in Grace’s medium containing 2 μM methoprene or the corresponding solvent (DMSO) alone. Importantly, methoprene treatment increased *Met, Kr-h1, Dsx,* and *Vg* mRNA levels by 1.5-, 149-, 1.9-, and 10-fold, respectively, (Fig. 6C). Moreoever, upon neck ligation, vitellogenesis and ovarian maturation were fully restored by methoprene treatment (Fig. 6D-F”). These data demonstrated that JH signaling alone is able to induce vitellogenesis and ovarian maturation even without additional stimulation by IIS and TOR.

### IIS and TOR promote vitellogenesis via inducing JH biosynthesis

To this end, we examined whether IIS and TOR promote vitellogenesis depending on or independent of JH biosynthesis using the neck-ligated cockroaches. Suprisingly, in contrast with methoprene treatment, treatments with bovine insulin (to restore IIS) or an amino acid mixture (to restore TOR) did not resuce ovarian maturation at all (Fig. 7A, B). As expected, western blot analyses showed that the levels of P-Akt and P-S6K in the fat body were restored to the control levels by treatments with bovine insulin and an amino acid mixture, respectively (Fig. 7C-D’). However, JH signaling and vitellogenesis were not restored at all by treatments with bovine insulin and an amino acid mixture (Fig. S5, 6). Finally, we determined whether IIS and TOR affect JH biosynthesis *in vitro.* CA incubation with LY294002 or rapamycin significantly decreased the expression of *Jhamt* and *Cyp 15a1* (Fig. 7E), in agreement with that IIS and TOR are required for the promotion of JH biosynthesis.

**Figure 7.**
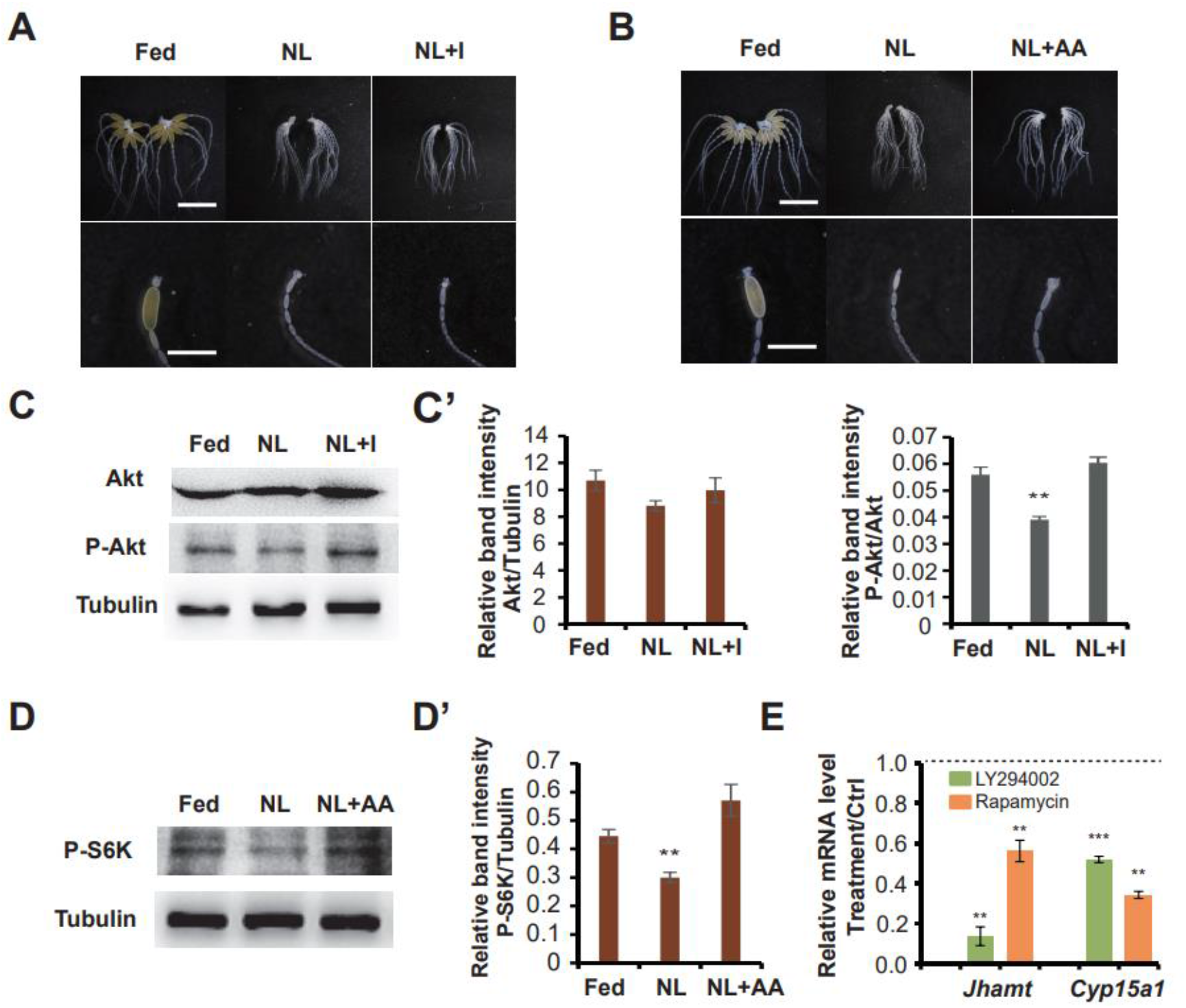
IIS and TOR promote vitellogenesis via inducing JH biosynthesis. **(A, B)** Morphological change of ovary after neck ligation (NL) and subsequent rescue with bovine insulin (NL+I) (A) and an amino acid mixture (NL+AA) (B). **(C, C’)** Changes of the protein levels of Akt and P-Akt after rescue with bovine insulin (C), quantification of band intensity showing the increase of P-Akt protein levels after bovine insulin treatment in the neck-ligated cockroaches (C’). **(D, D’)** Changes of the protein levels of P-S6K after rescue with an amino acid mixture (D), quantification of band intensity showing the increase of P-S6K protein levels after an amino acid mixture treatment in the neck-ligated cockroaches (D’). **(E)** Expression of *Jhamt* and *Cyp15a1* in the *in vitro* cultured CA treated with LY294002 or rapamycin. \*\**P*<0.01, \*\*\**P*<0.001, compared to the negative control (DMSO). n=4.

In summary, IIS and TOR do not induce vitellogenesis independent of JH in the American cockroach, they promote vitellogenesis in an indirect manner via activating JH biosynthesis (Fig. 8).

**Figure 8.**
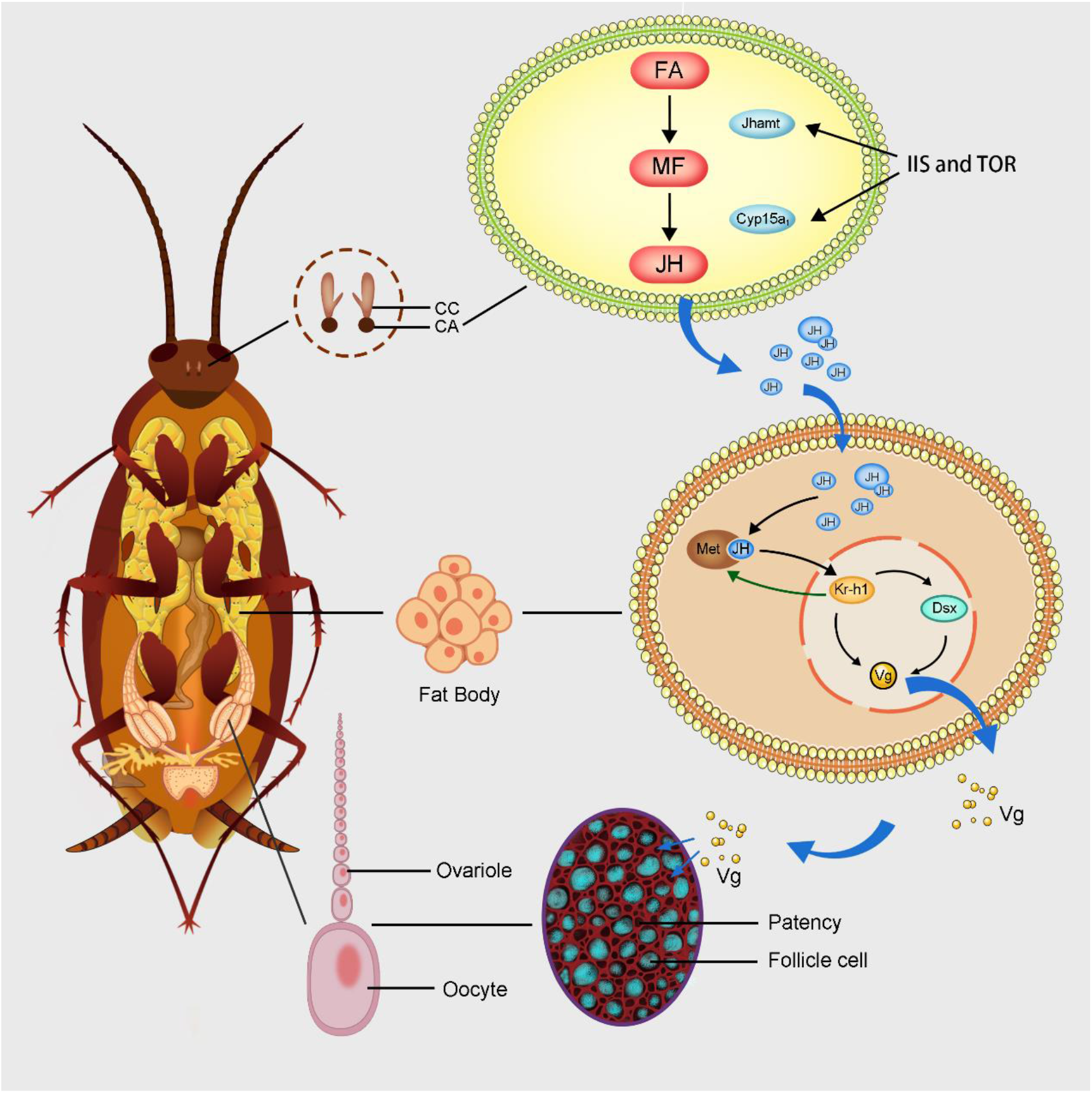
Schematic view of IIS and TOR promote vitellogenesis via inducing JH biosynthesis in the American cockroach. Upon feeding during the first reproductive cycle, IIS and TOR was activiated to induce the expression of *Jhamt* and *Cyp15a1* in the CA and thus to activate JH biosynthesis. Via the genomic action, JH functions through Met and Kr-h1, which form a positive feedback regulatory loop, to induce *Vg* expression in the fat body partially via Dsx. Meanwhile, via the nongenomic action, JH induces the development of functional follicular patency to allow the internalization of hemolymph Vg into maturing oocytes. Moreover, the JH-stimulated Vg production is a prerequisite for the JH-activated follicular patency and thus Vg uptake highlighting the coordiative actions of JH in the regulation of vitellogenesis.

## DISCUSSION

### IIS and TOR promote vitellogenesis by inducing JH biosynthesis

The interplay between the nutrient sensors (IIS and TOR) and JH in the regulation of insect vitellogenesis varies depending on their reproductive strategies. In the mosquito *Aedes aegypti,* IIS and TOR stimulate the biosynthesis of JH, which controls a critical previtellogenic preparatory phase in the fat body but not Vg production (Perez-Hedo et al., 2013; Roy et al., 2018). Later in the vitellogenesis phase, IIS and TOR coordinate with 20-hydroxyecdysone to induce *Vg* expression in the fat body and ovarian maturation (Hansen et al., 2004; Roy et al., 2007). JH might also affect insulin production to regulate reproduction in the honeybee *Apis mellifera* (Corona et al., 2007) and *T. castaneum* (Sheng et al., 2011). Nevertheless, in our preliminary experiments, no effects of JH on insulin production were observed in the American cockroach. A number of studies have shown that IIS and TOR promote both JH biosynthesis and Vg production in the German cockroach *Blattella germanica* (Abrisqueta et al., 2014; Maestro et al., 2009) and *T. castaneum* (Parthasarathy and Palli, 2011; Parthasarathy et al., 2010), but the interplay between the nutrient sensors and JH has not been fully demonstrated yet.

In the American cockroach, we have also determined that IIS and TOR promote both JH biosynthesis and vitellogenesis (Fig. 2–5). More importantly, we demonstrated that IIS and TOR promote vitellogenesis in an indirect manner by activating JH biosynthesis, and that IIS and TOR alone do not promote vitellogenesis (Fig. 6, 7). Neck ligation is a classic physiology procedure that was wildly used for functional studies by insect physiologists in the past. Neck ligation not only prevented the cockroaches from feeding, but also depleted the production of both insulin and JH in the brain. Thus, in the neck-ligated (both nutrient- and JH-deficient) cockroaches, both JH biosynthesis and vitellogenesis were completely abolished. Crucially, JH, but not IIS and TOR activitly, restored vitellogenesis in the neck-ligated cockroaches. Therefore, combining both classic physiology (i.e. neck ligation and microinjection) and morden molecular techniques (i.e. RNAi), we revealed the complex interplay between the nutrient sensors and JH in the regulation of vitellogenesis in the Amercian cockroach. We suppose this model (Fig. 8) applies to the a number of insect groups, such as cockroaches, locusts, and beetles, in which JH is the major hormone controlling vitellogeneis. In this study, we not only provided a conclusive model of endocrinal and nutritional regulation of vitellogenesis in the American cockroach (Fig. 8), but also refreshed the classic physiology procedure (neck ligation) as a promising method to study the interactions of signal transduction pathways in insects.

### JH signaling and its regulation of vitellogenesis

As discussed above, in this study, we have concluvisely determined that JH induces vitellogenesis even in the absence of additional stimulation by IIS or TOR (Fig. 2, 3, 6). Interestingly, we observed that Met, the JH receptor, responded to JH at the transcription level (Fig. 6B-C) and that Kr-h1, which is downstream of Met, may also have a positive feedback regulatory effect on *Met* expression (Fig. 3C), similar to the reports in the plant brown hopper *Nilaparvata lugens* (Lin et al., 2015). It is likely that the positive feedback regulatory loop between Met and Kr-h1 might maximize the JH signaling in the regulation of vitellogenesis.

Based on the previous studies (Burtis et al., 1991; Gotoh et al., 2014; Shukla and Nagaraju, 2010; Shukla and Palli, 2012), we hypothesize that Dsx might be a molecular link in the JH regulation of vitellogenesis. In this study, we found that JH induced *Dsx* expression and Dsx is required for JH signaling to promte vitellogenesis, establishing Dsx as a molecular connection between JH signaling and vitellogenesis (Fig. 2C, 6B-C). However, RNAi depetion of *Dsx* did not completely ablolish JH-induced *Vg* expression in the fat body (Fig. 2, S2), suggesting that some other transcriptional factors (i.e. GATA) (Park et al., 2006) should be involved in the JH induction of *Vg* expression and thus vitellogenesis. In addition, we are currently examining if JH stimulates endocycle and DNA replication in the fat body to prepare for Vg production, as deomonstrated in the locust *Locusta migratoria* (Guo et al., 2014; Luo et al., 2017; Wu et al., 2016, 2018).

Our data also support that the nongenomic action of JH might promote the development of functional follicular patency in the maturing oocytes and thus Vg uptake (Davey, 2000; Bai and Palli, 2016). In agreement with the previous review reports (Jindra et al., 2013; Li et al., 2019; Roy et al., 2018), we suppose that JH promotes vitellogenesis by multiple ways in the American cockroach, including the promotion of Vg production in the fat body via the genomic action as well as the development of follicular patency and thus Vg uptake in the maturing oocytes via the non-genomic action. Moreover, we have revealed that Vg production might alter ovarian maturation by affecting the follicular patency and thus Vg uptake (Fig. 1C-D’), strengthening the coordiatiom of genomic and nongenomic actions of JH in the regulation of vitellogenesis.

In summary, our composite data have revealed the molecular mechanisms underlying the endocrinal and nutritional regulation of vitellogenesis in the Amercian cockroach. Upon feeding during the first reproductive cycle, IIS and TOR was activiated to induce the expression of *Jhamt* and *Cyp15a1* in the CA and thus to activate JH biosynthesis. Via the genomic action, JH functions through Met and Kr-h1, which form a positive feedback regulatory loop, to induce *Vg* expression in the fat body partially via Dsx. Meanwhile, via the nongenomic action, JH induces the development of functional follicular patency to allow the internalization of hemolymph Vg into maturing oocytes. Moreover, the JH-stimulated Vg production is a prerequisite for the JH-activated follicular patency and thus Vg uptake, highlighting the coordiative actions of JH in the regulation of vitellogenesis. Combining classic physiology with modern molecular techniques, we have demonstrated that IIS and TOR promote vitellogenesis via inducing JH biosynthesis in the American cockroach (Fig. 8).

## MATERIALS AND METHODS

### Animals

The cockroaches used in the present study have been previously described (Li et al., 2018). The animals were kept at 28-30°C under a 12/12 h light/dark cycle and relative humidity of 70–80% in plastic cages and fed commercial rat food and water. To obtain pools of synchronized animals, newly emerged female adults were picked from the colony and placed in separate containers. Operations were performed after anaesthetization with CO2. Dissection and tissue sampling were carried out in cockroach saline solution. For all experiments described in this paper, at least three biological repeats were performed with 30 animals in each replicate.

### RNAi

For RNAi, 300–500 bp fragments of the target genes were PCR amplified from the complementary DNA. Then, primers attached to the T7 promoter sequence were used for the PCR amplification of double-stranded RNA (dsRNA) templates. The dsRNA was synthesized using a T7 RiboMAX Express RNAi kit (Promega, USA) according to the manufacturer’s instructions. A 92 bp noncoding sequence from the pSTBlue-1 vector (CK) was used as a control dsRNA (Gomez-Orte and Belles, 2009). The sequences of all primers used for dsRNA synthesis in this study are listed in Table S1. dsRNA was quantified using a Nanodrop spectrophotometer (Thermo Scientific, USA), and 3 μg of dsRNA was injected into the abdomen of each animal. The injection was carried out on day 2, 4, and 6 PAE (more details in Fig. S7A). The control animals were injected with the same dose of CK dsRNA at the same time.

### LY294002 and rapamycin application

Cockroaches were injected with 4 μg of LY294002 (a PI3K inhibitor) (MCE, USA) or rapamycin (MCE, USA) on day 2 and day 4 PAE. Control animals were injected with the corresponding volume of the solvent. Tissues were collected at day 5 PAE for further use (more details in Fig. S7B).

### Methoprene, bovine insulin, and amino acid mixture application

The females of day 2 PAE were ligated with dental floss at the neck. A total of 100 μg of methoprene (MCE, USA) in 2 μl of DMSO (JH analog application), 25 μg of bovine insulin (Yuanye Bio-Technology, China) in 2 μl of 1 M acetic acid or 2 μl of a standard solution of an amino acid mixture (type H) (Wako, Japan) was injected into the abdomen, respectively. Control animals were injected with the corresponding volume of the solvent. Ovaries were collected 72 h after injection for imaging, follicular epithelium staining and protein extraction (more details in Fig. S7C).

### Tissue imaging and confocal microscopy

Cockroaches were dissected in CSS under an Olympus SZ61 microscope (Japan). To observe the ovary developmental pattern, the ovaries of adult females from day 1 to day 8 PAE were dissected. For the RNAi experiment, the ovaries were dissected on day 5 or day 7 PAE; body weight, ovarian weight, and primary oocyte length were recorded at the same time. Images of the ovaries and ovarioles were captured with a Nikon DS-Ri2 camera and Nikon SMZ25 microscope (Japan). Primary oocyte length was measured using NIS-Elements BR 4.50.00 software. For cell staining, ovarioles were fixed in 4% paraformaldehyde for 60 min and permeabilized in 0.1% Triton X-100 for an additional 15 min. F-Actin was stained with TRITC-phalloidin (excitation wavelength 545 nm) (Yeasen, China). Nuclei were stained with Hoechst 33342 (excitation wavelength 350 nm) (Yeasen, China). The images were captured with an Olympus FluoView FV3000 confocal microscopy and analyzed with FV31S-SW software (Olympus).

### Tissue incubation *in vitro*

Fat bodies adhered to abdominal tergites and epidermises were dissected from freshly emerged adult females or females on 1 or 5 days PAE. The CA was dissected from adult females on day 5 PAE. The tissue or organ was then preincubated for 30 min in 1 ml of Grace’s insect medium (Thermo Scientific, USA) at 30°C in the dark (Maestro et al., 2009). After preincubation, fat bodies from females on day 1 PAE were incubated for 6 h in medium supplemented with 2 μM methoprene. Fat bodies or CA from females on day 5 PAE was incubated for 6 h in medium supplemented with 2 μM methoprene or 50 mM LY294002 or rapamycin. Controls were incubated with the corresponding volume of the solvent. After the final incubation, tissues were frozen in liquid nitrogen immediately and stored at −80°C until RNA extraction.

### Total RNA extraction and qPCR

Twenty-four hours after the injection of dsRNA or reagents, abdominal fat body tissues and the heads of the cockroaches were collected, flash-frozen in liquid nitrogen to prevent RNA degradation and stored at −80°C until further processing. Total RNA was extracted from the fat body, head, or CA using a Direct-zolTM RNA MiniPrep (Zymo Research, USA) according to the manufacturer’s instructions. Reverse transcription was performed by using reverse transcriptase M-MLV (TaKaRa, Japan) according to the manufacturer’s instructions. qPCR was performed using SYBR^®^ Select Master Mix (Applied Biosystems, USA) and the Applied Biosystems™ QuantStudio™ 6 Flex Real-Time PCR System. *Actin* was chosen as a reference gene for qPCR analysis. The sequences of all primers used for qPCR in this study are listed in Table S2.

### Western blots

Total proteins were extracted from the fat body of adult females. Tissue lysates in RIPA lysis buffer (Beyotime Biotechnology, China) with 1 mM PMSF and a phosphatase inhibitor cocktail (CWBIO, China) were then cleared by centrifugation at 12,000 × g at 4°C for 30 min. Extracted proteins were quantified using a BCA protein assay kit (Yeasen, China), fractionated with 10% SDS-PAGE, and then transferred to PVDF membranes (Millipore, USA). Western blotingt was performed using antibodies against Akt, P-Akt, and P-S6K (Cell Signaling Technology, USA). Tubulin (Beyotime Biotechnology, China) was used as a loading control. Bands were imaged with a Tanon 5200 system (Tanon, China) and analyzed using ImageJ software.

## Data analysis

Statistical analyses were performed with Student’s *t*-test or one-way ANOVA. Values are shown as the mean ± standard deviation (SD), and differences were considered significant at *P* < 0.05.

## Acknowledgments

We thank Drs. Lynn Riddiford and Marek Jindra for advice and comments on improving the manuscript.

## Author contributions

Conceptualization: S.L.; Methodology: S.Z., F.L., C.R., Y.Q., S.L.; Software: S.Z., F.L., Y.Q., S.L.; Formal analysis: S.Z., F.L., Y.Q., S.L.; Investigation: S.Z.; F.L.; H.Z.; N. L.; C.R., Y.S., S.Z.; S.P.; J.W.; Y.Q.; S.L.; Resources: S.Z., F.L., Y.Q., S.L.; Data curation: S.Z., F.L., Y.Q., S.L.; Writing – original draft: S.Z., F.L.; Writing – review & editing: S.Z.; F.L.; C.R., Y.S., S.Z.; S.P.; G.W., J.W.; Y.Q.; S.L.; Supervision: Y.Q., S.L.; Project administration: S.Z., S.L.; Funding acquisition: N.L.; C.R.; J.W., Y.Q.; S.L.

## Supplementary Figures

**Fig. S1.**
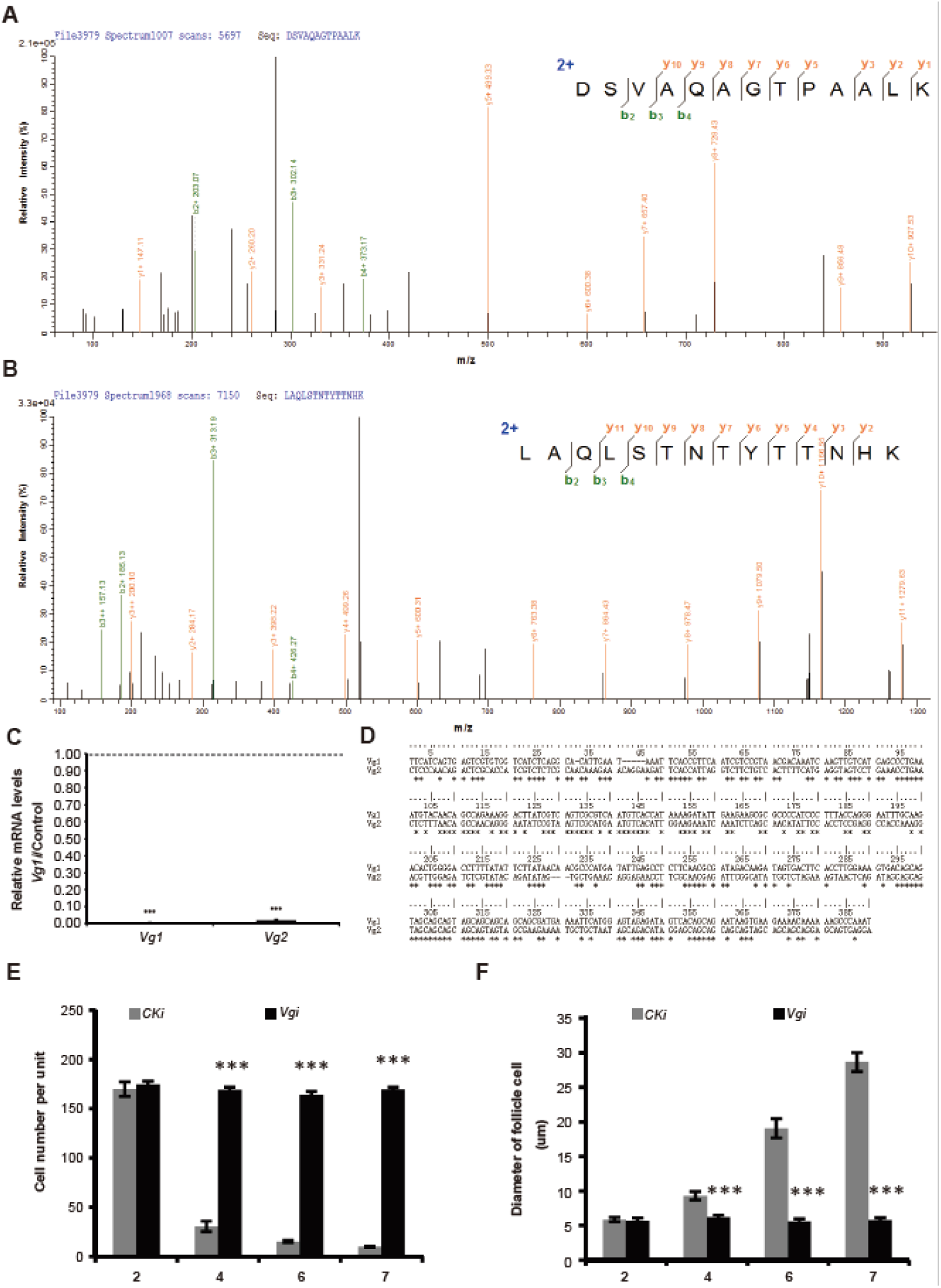
Identification of Vg by mass spectroscopy and the effect of *Vg1* RNAi on the follicular epithelium. **(A, B)** The identification of Vg proteins by mass spectrometry (the 100 kDa band). Total proteins were extracted from the ovaries. n=3. (A) The DSVAQAGTPAALK peptide fragment of Vg1 protein detected in the ovary proteins; (B) The LAQLSTNTYTTNHK peptide fragment of Vg2 detected in the ovary proteins. **(C)** Effect of *Vg*1 RNAi on the expression of *Vg*1 and *Vg*2. *(D)* Alignment of the DNA fragment of *Vg*1 used to design *Vg*1 dsRNA and the related fragment of *Vg*2.. **(E, F)** Effects of *Vg* RNAi on follicle cell number (E) and nuclei diameter (F). \*\*\**P*<0.001, compared to the negative control (CK RNAi). n=3.

**Fig. S2.**
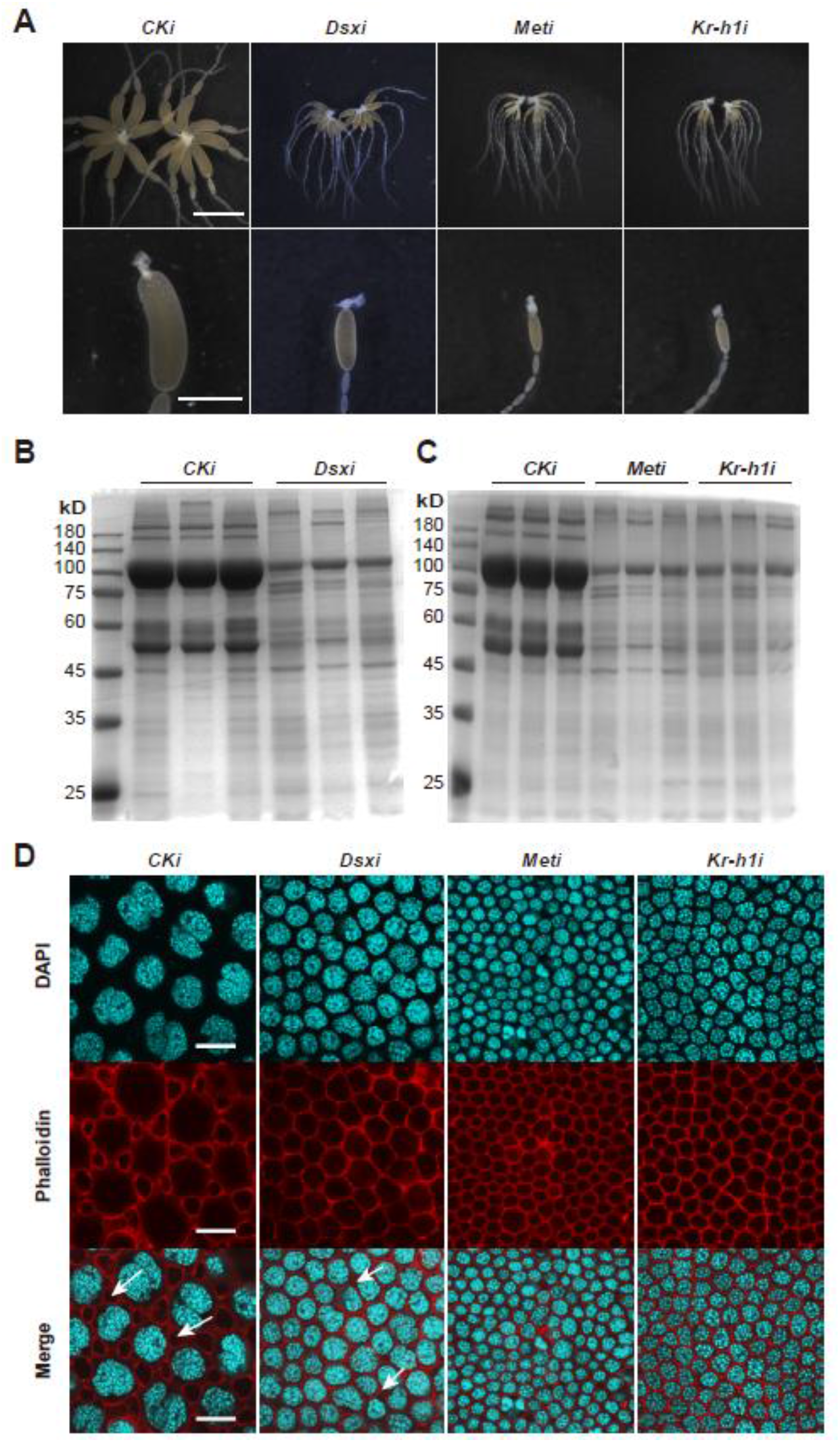
Depletion of JH signaling inhibited ovarian maturation. **(A)** Representative phenotypes of ovaries and primary oocytes after *Dsx* RNAi, *Met* RNAi, or *Kr-h1* RNAi versus the negative control (CK). Scale bar: 5mm (upper panels), 2mm (lower panels). n=10. **(B)** Effect of *Dsx* RNAi on total ovary protein versus the negative control (CK). n=3. **(C)** Effect of *Met* RNAi and *Kr-h1* RNAi on total ovary protein versus the negative control (CK). n=3. **(D)** Effect of *Dsx* RNAi, Met RNAi and Kr-h1 RNAi on the follicular epithelium. Blue, follicle cell nuclei; red, F-actin. White arrows indicate patency. Scale bar: 20 μm. n=3.

**Fig. S3.**
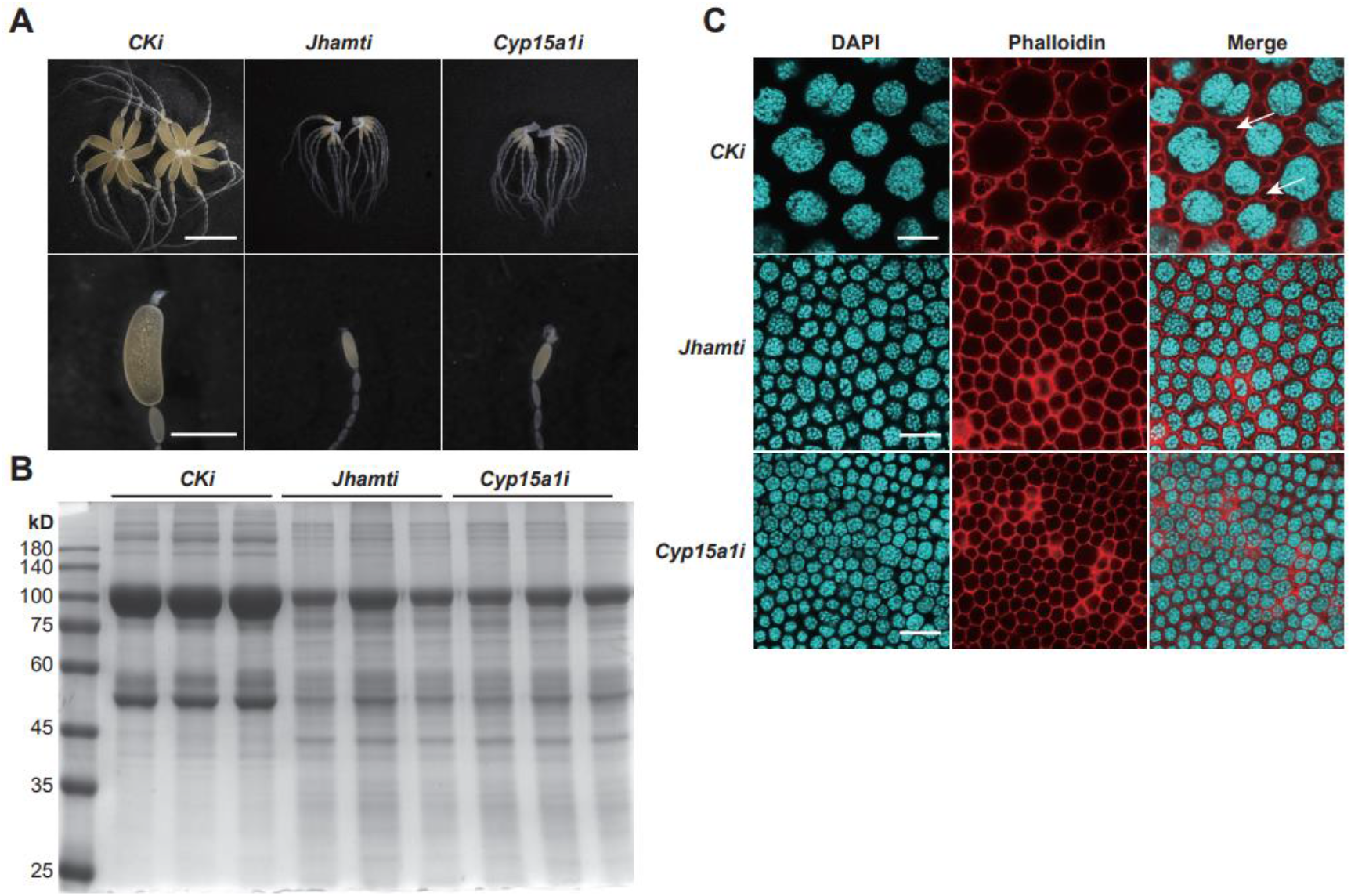
Depletion of JH biosynthetic pathway inhibited ovarian maturation. **(A)** Representative phenotypes of ovaries and primary oocytes after *Jhamt* RNAi and *Cyp15a1* RNAi versus the negative control (CK). Scale bar: 5mm (upper panels), 2mm (lower panels). n=10. **(B)** Effect of *Jhamt* RNAi and *Cyp15a1* RNAi on total ovary protein versus the negative control (CK). n=3. **(C)** Effect of *Jhamt* RNAi and *Cyp15a1* RNAi on the follicular epithelium. Blue, follicle cell nuclei; red, F-actin. White arrows indicate patency. Scale bar: 20 μm. n=3.

**Fig. S4.**
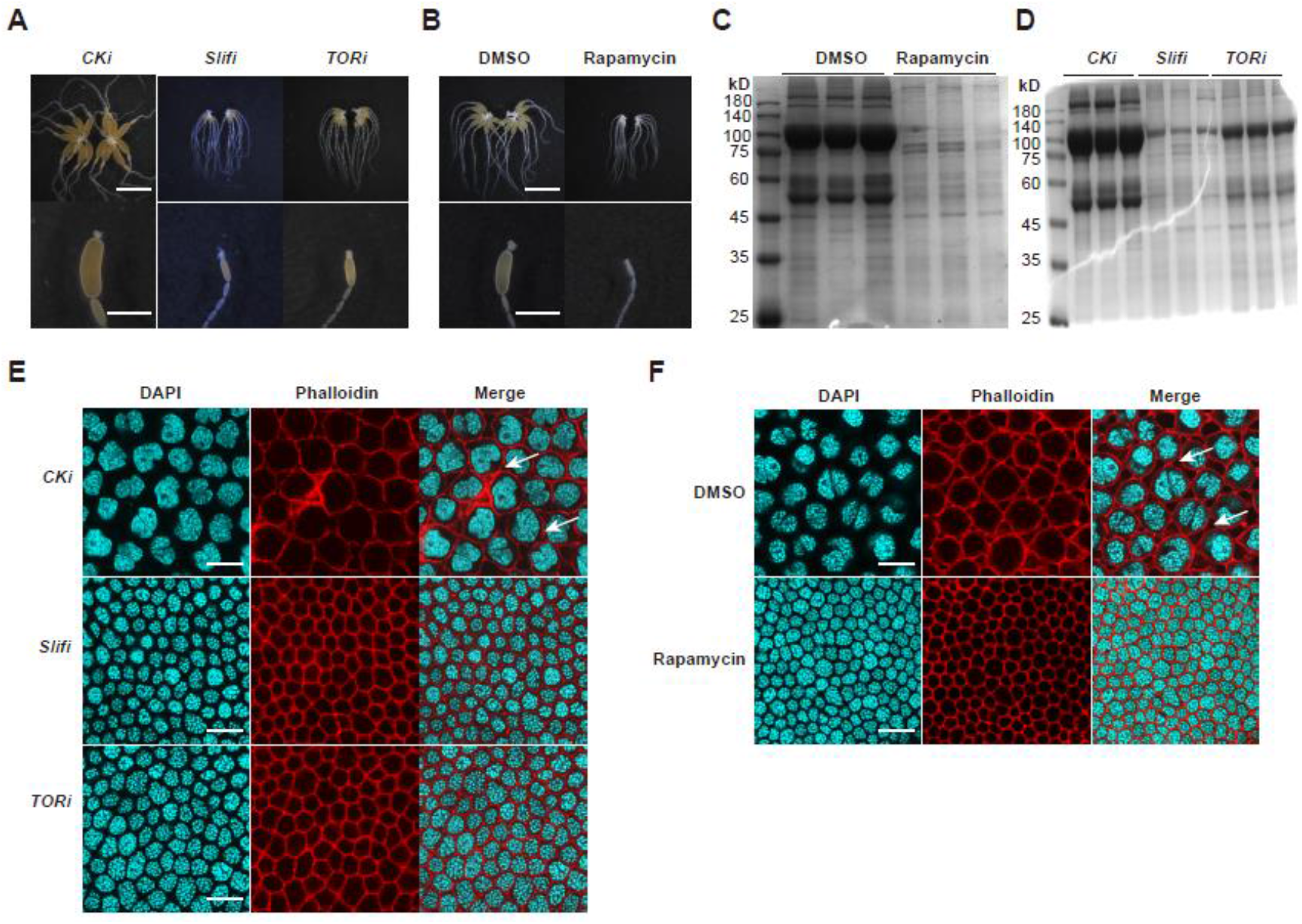
Depletion of TOR inhibited ovarian maturation. **(A, B)** Representative phenotypes of ovaries and primary oocytes after (A) *Slif* RNAi, *TOR* RNAi, or (B) rapamycin treatment. Scale bar: 5mm (upper panels), 2mm (lower). n=10. **(C, D)** Effect of *Slif* RNAi, *TOR* RNAi, or rapamycin treatment on total ovary protein compared to the negative control (CK RNAi or DMSO). n=3. **(E, F)** Effect of *Slif* RNAi, *TOR* RNAi, or rapamycin treatment on the follicular epithelium. Blue, follicle cell nuclei; red, Factin. White arrows indicate patency. Scale bar: 20 μm.

**Fig. S5.**
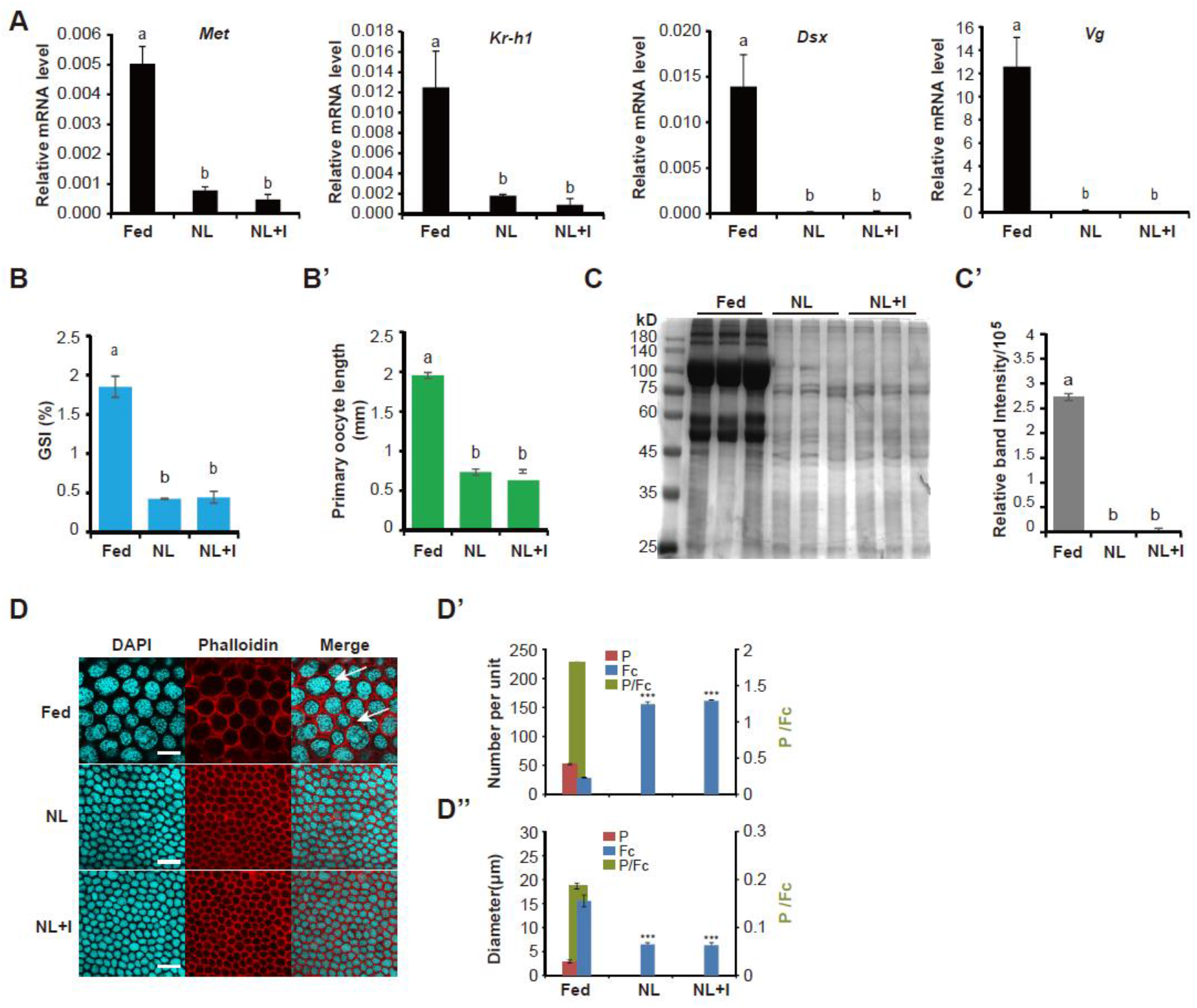
Bovine insulin did not stimulate vitellogenesis in the absemce of JH. Effects of neck ligation (NL) and subsequent rescue with bovine insulin (NL+I) on the expression of *Vg*, *Dsx*, *Met*, and *Kr-h1* in the fat body (A), the gonadosomatic index (GSI) and the primary oocyte length (B, B’), the 100 kDa band intensity (C, C’), number of Fc and P and the number index (P/Fc), diameter of Fc nuclei and P and the diameter index (P/Fc) (D-D”). Blue, follicle cell nuclei; red, F-actin. n=3. White arrows indicate patency. Scale bar: 20 μm. \*\*\**P*<0.001. The bars labeled with different letters indicate significant difference at *P*< 0.05.

**Fig. S6.**
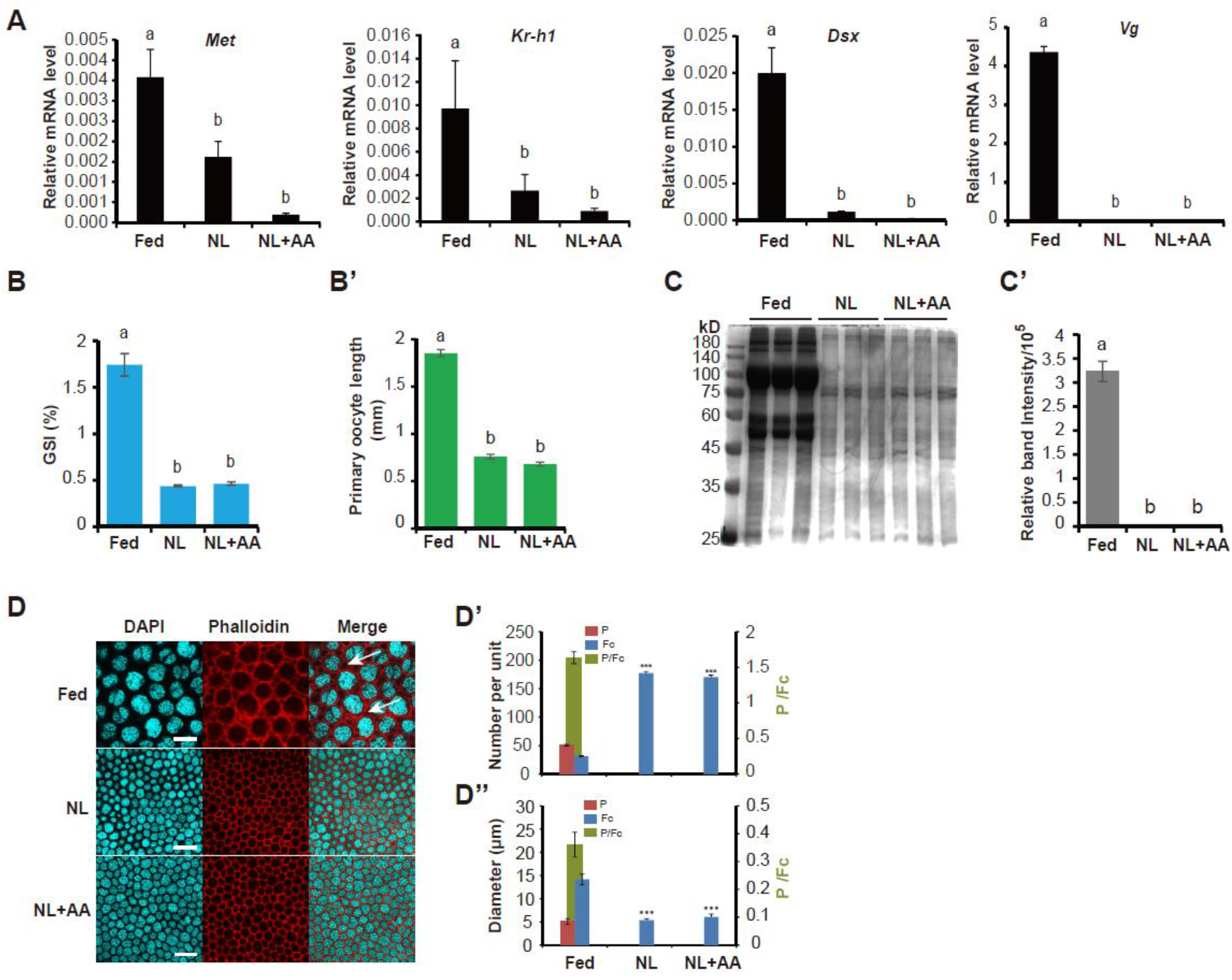
Amino acid did not stimulate vitellogenesis independent of JH. Effects of neck ligation (NL) and subsequent rescue with an amino acid mixture (NL+AA) on the expression of *Vg*, *Dsx*, *Met*, and *Kr-h1* in the fat body (A), the gonadosomatic index (GSI) and the primary oocyte length (B, B’), the 100 kDa band intensity (C, C’), number of Fc and P and the number index (P/Fc), diameter of Fc nuclei and P and the diameter index (P/Fc) (D-D”). Blue, follicle cell nuclei; red, F-actin. n=3. White arrows indicate patency. Scale bar: 20 μm. \*\*\**P*<0.001. The bars labeled with different letters indicate significant difference at *P*< 0.05.

**Fig. S7.**
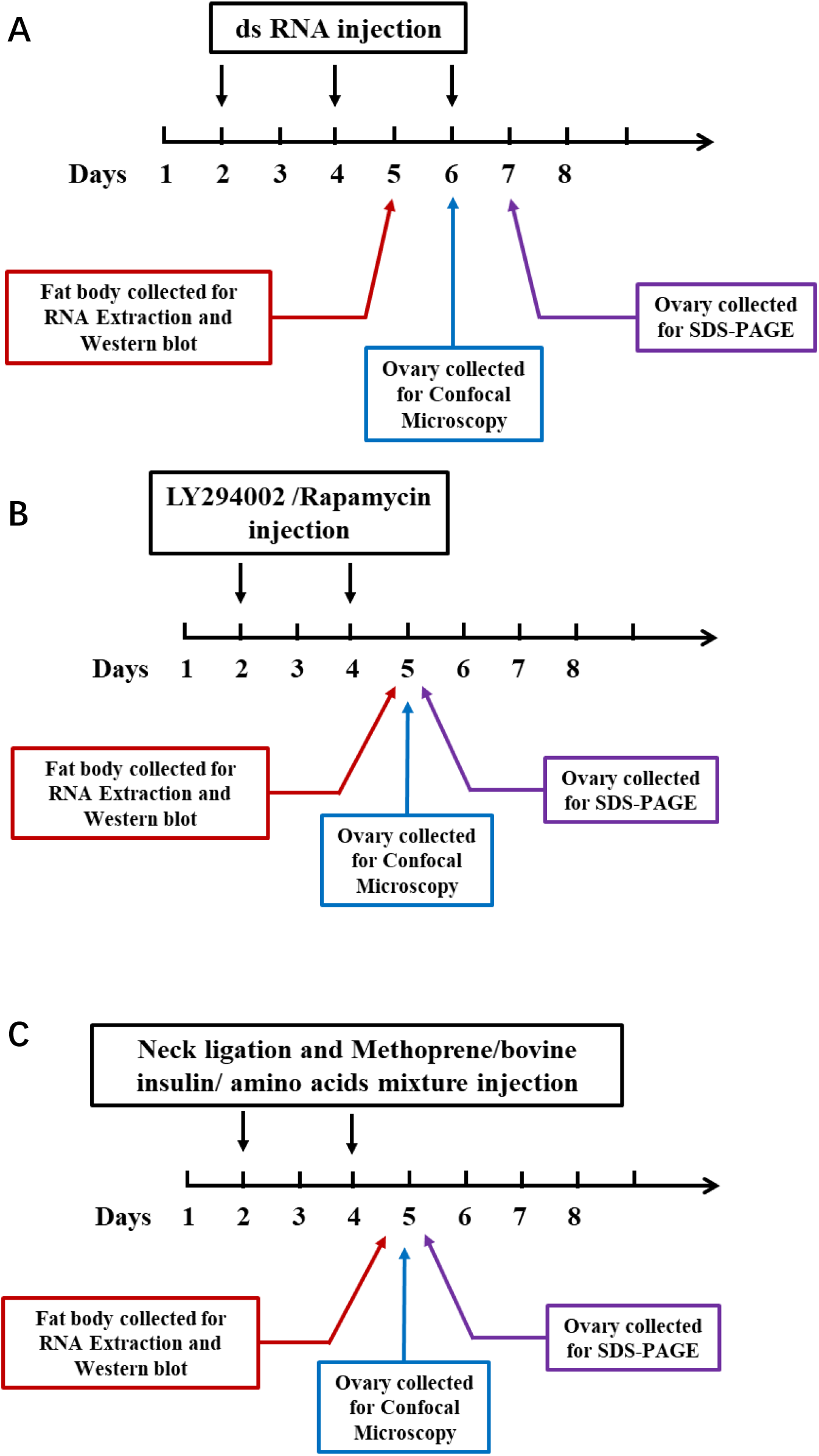
Strategy for experiments. **(A)** Strategy of RNAi. **(B)** Strategy of depletion of IIS and TOR by inhibitors. **(C)** Strategy for rescue experiments after physically blocking JH signaling and IIS/TOR.

**Table S1.**
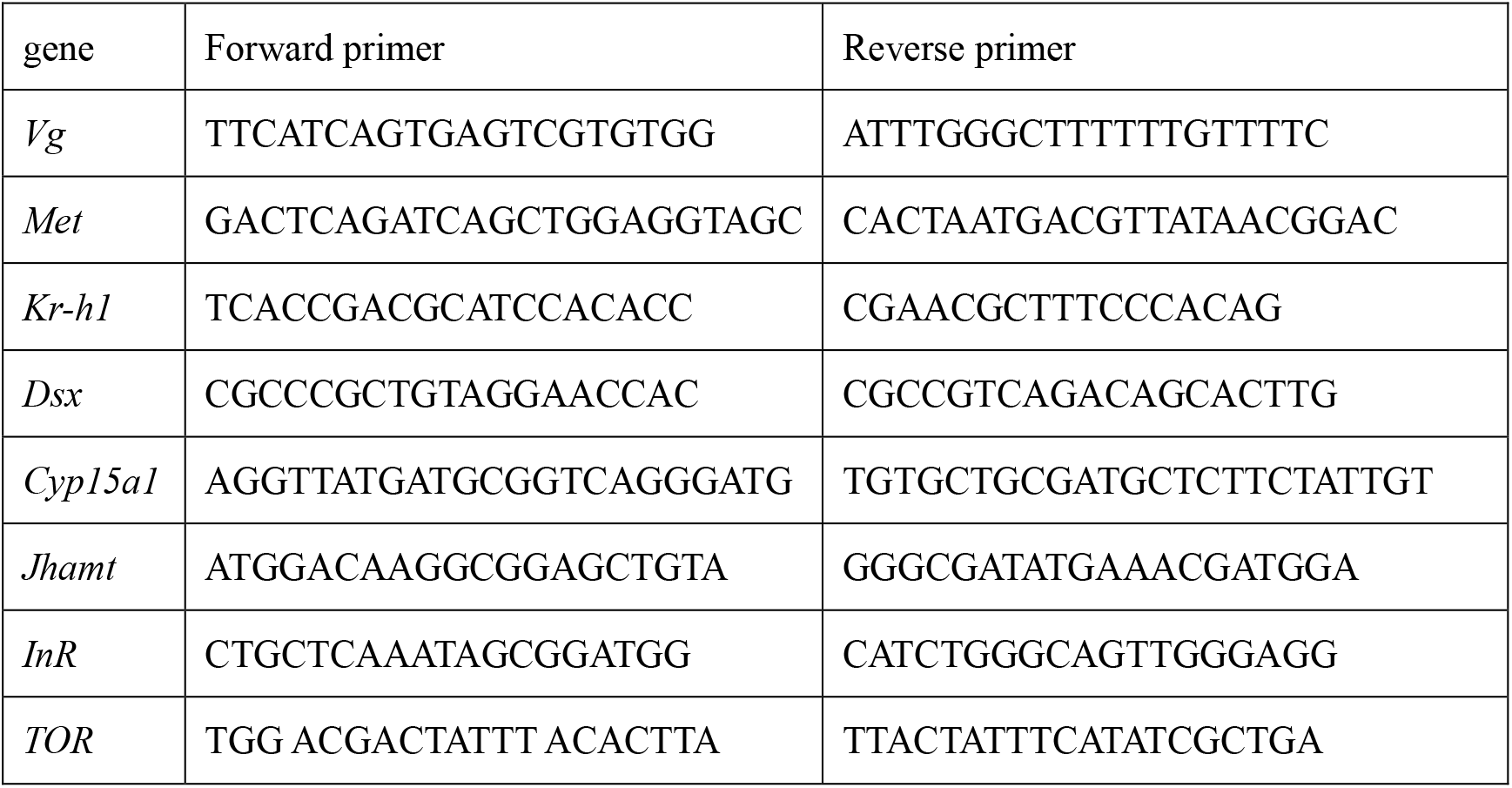
PCR primers for dsRNA.

**Table S2.**
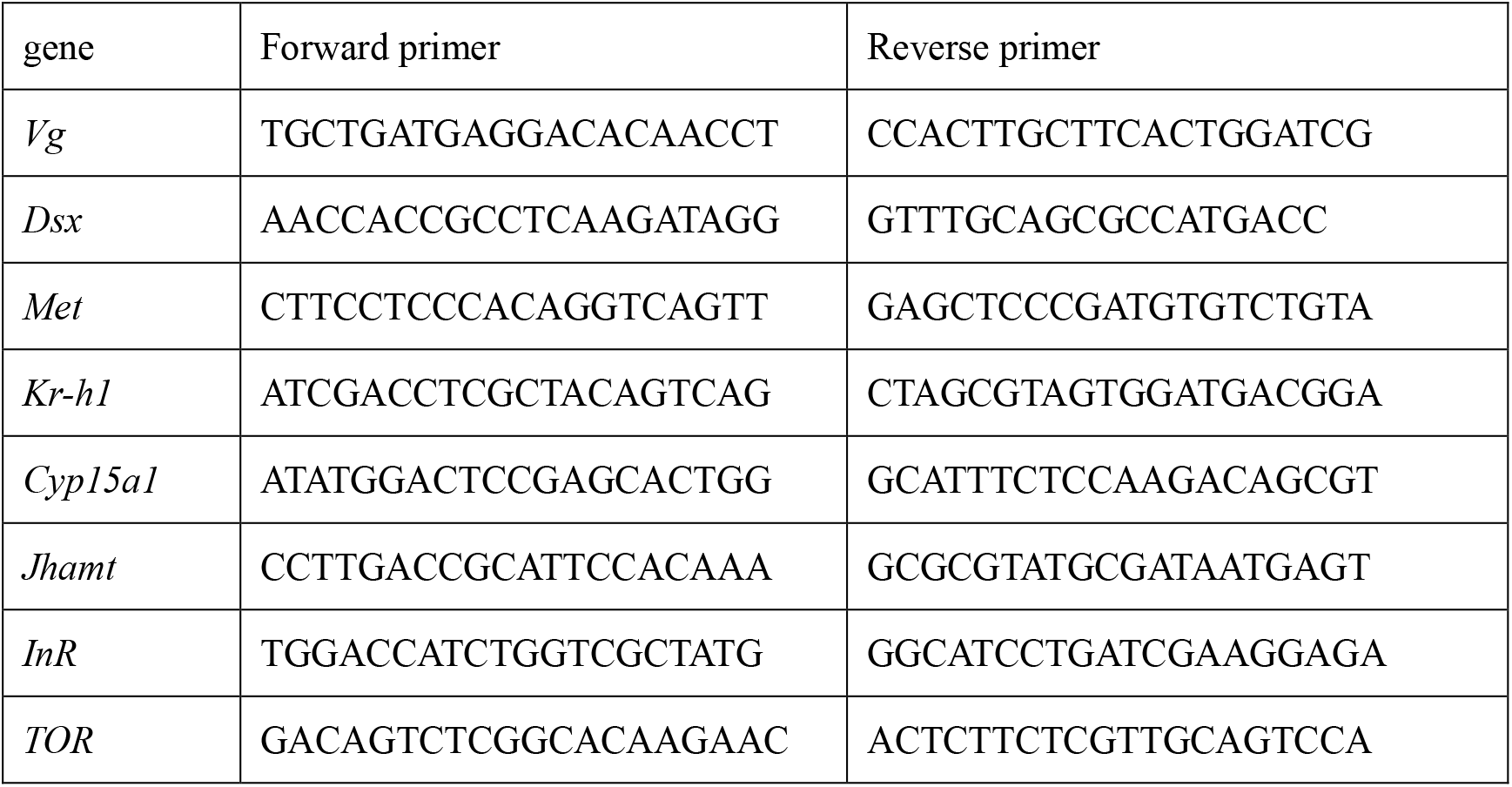
Primers for qPCR

## References

Abrisqueta, M., Suren-Castillo, S. and Maestro, J. L. (2014). Insulin receptor-mediated nutritional signalling regulates juvenile hormone biosynthesis and vitellogenin production in the German cockroach. Insect Biochem Mol Biol 49, 14–23. doi: 10.1016/j.ibmb.2014.03.005

Bai, H. and Palli, S. R. (2016). Identification of G protein-coupled receptors required for vitellogenin uptake into the oocytes of the red flour beetle, *Tribolium castaneum*. Sci Rep 6, doi: ARTN 27648 10.1038/srep27648

Belles, X., Martin, D. and Piulachs, M. D. (2005). The mevalonate pathway and the synthesis of juvenile hormone in insects. Annu Rev Entomol 50, 181–199. doi: 10.1146/annurev.ento.50.071803.130356

Burtis, K. C., Coschigano, K. T., Baker, B. S. and Wensink, P. C. (1991). The double sex proteins of Drosophila melanogaster bind directly to a sex-specific yolk protein gene enhancer. EMBO J 10, 2577–2582. doi: 10.1002/j.1460-2075.1991.tb07798.x

Colombani, J., Raisin, S., Pantalacci, S., Radimerski, T., Montagne, J. and Leopold, P. (2003). A nutrient sensor mechanism controls Drosophila growth. Cell 114, 739–749. doi: 10.1016/s0092-8674(03)00713-x

Corona, M., Velarde, R. A., Remolina, S., Moran-Lauter, A., Wang, Y., Hughes, K. A. and Robinson, G. E. (2007). Vitellogenin, juvenile hormone, insulin signaling, and queen honey bee longevity. Proc Natl Acad Sci U S A 104, 7128–7133. doi: 10.1073/pnas.0701909104

Davey, K. G. (2000). The modes of action of juvenile hormones: some questions we ought to ask. Insect Biochem Mol Biol 30, 663–669. doi: 10.1016/S0965-1748(00)00037-0

Defelipe, L. A., Dolghih, E., Roitberg, A. E., Nouzova, M., Mayoral, J. G., Noriega, F. G. and Turjanski, A. G. (2011). Juvenile hormone synthesis: “esterify then epoxidize” or “epoxidize then esterify”? Insights from the structural characterization of juvenile hormone acid methyltransferase. Insect Biochem Mol Biol 41, 228–235. doi: 10.1016/j.ibmb.2010.12.008

Gomez-Orte, E. and Belles, X. (2009). MicroRNA-dependent metamorphosis in hemimetabolan insects. P Natl Acad Sci USA 106, 21678–21682. doi: 10.1073/pnas.0907391106

Gotoh, H., Miyakawa, H., Ishikawa, A., Ishikawa, Y., Sugime, Y., Emlen, D. J., Lavine, L. C. and Miura, T. (2014). Developmental Link between Sex and Nutrition; doublesex Regulates Sex-Specific Mandible Growth via Juvenile Hormone Signaling in Stag Beetles. Plos Genet 10, doi: ARTN e1004098 10.1371/journal.pgen.1004098

Hansen, I. A., Attardo, G. M., Park, J. H., Peng, Q. and Raikhel, A. S. (2004). Target of rapamycin-mediated amino acid signaling in mosquito anautogeny. P Natl Acad Sci USA 101, 10626–10631. doi: 10.1073/pnas.0403460101

Jindra, M., Palli, S. R. and Riddiford, L. M. (2013). The Juvenile Hormone Signaling Pathway in Insect Development. Annu Rev Entomol, 58, 181–204. doi: 10.1146/annurev-ento-120811-153700

Jing, Y. P., An, H. L., Zhang, S. J., Wang, N. B. and Zhou, S. T. (2018). Protein kinase C mediates juvenile hormone-dependent phosphorylation of Na+/K+-ATPase to induce ovarian follicular patency for yolk protein uptake. J Bio Chem 293, 20112–20122. doi: 10.1074/jbc.RA118.005692

Li, S., Yu, X. Q. and Feng, Q. L. (2019). Fat Body Biology in the Last Decade. Annu Rev Entomol 64, 315–333. doi: 10.1146/annurev-ento-011118-112007

Li, S., Zhu, S. M., Jia, Q. Q., Yuan, D. W., Ren, C. H., Li, K., Liu, S. N., Cui, Y. Y., Zhao, H. G., Cao, Y. H., et al. (2018). The genomic and functional landscapes of developmental plasticity in the American cockroach. Nat Commun 9, doi: ARTN 1008 10.1038/s41467-018-03281-1

Lin, X. D., Yao, Y. and Wang, B. (2015). Methoprene-tolerant (Met) and Krupple-homologue 1 (Kr-h1) are required for ovariole development and egg maturation in the brown plant hopper. Sci Rep 5, doi: ARTN 18064 10.1038/srep18064

Liu, P. C., Peng, H. J. and Zhu, J. S. (2015). Juvenile hormone-activated phospholipase C pathway enhances transcriptional activation by the methoprene-tolerant protein. Proc Natl Acad Sci USA 112, E1871–E1879. doi: 10.1073/pnas.1423204112

Luo, M. W., Li, D., Wang, Z. M., Guo, W., Kang, L. and Zhou, S. T. (2017). Juvenile hormone differentially regulates two Grp78 genes encoding protein chaperones required for insect fat body cell homeostasis and vitellogenesis. J Bio Chem 292, 8823–8834. doi: 10.1074/jbc.M117.780957

Maestro, J. L., Cobo, J. and Belles, X. (2009). Target of Rapamycin (TOR) Mediates the Transduction of Nutritional Signals into Juvenile Hormone Production. J Biol Chem 284, 5506–5513. doi: 10.1074/jbc.M807042200

Nouzova, M., Edwards, M. J., Mayoral, J. G. and Noriega, F. G. (2011). A coordinated expression of biosynthetic enzymes controls the flux of juvenile hormone precursors in the corpora allata of mosquitoes. Insect Biochem Mol Biol 41, 660–669. doi: 10.1016/j.ibmb.2011.04.008

Ojani, R., Liu, P. C., Fu, X. N. and Zhu, J. S. (2016). Protein kinase C modulates transcriptional activation by the juvenile hormone receptor methoprene-tolerant. Insect Biochem Mol Biol 70, 44–52. doi: 10.1016/j.ibmb.2015.12.001

Park, J. H., Attardo, G. M., Hansen, I. A. and Raikhel, A. S. (2006). GATA factor translation is the final downstream step in the amino acid/target-of-rapamycin-mediated vitellogenin gene expression in the anautogenous mosquito *Aedes aegypti*. J Bio Chem 281, 11167–11176. doi: 10.1074/jbc.M601517200

Parthasarathy, R. and Palli, S. R. (2011). Molecular analysis of nutritional and hormonal regulation of female reproduction in the red flour beetle, *Tribolium castaneum*. Insect Biochem Mol Biol 41, 294–305. doi: 10.1016/j.ibmb.2011.01.006

Parthasarathy, R., Sun, Z. Y., Bai, H. and Palli, S. R. (2010). Juvenile hormone regulation of vitellogenin synthesis in the red flour beetle, *Tribolium castaneum*. Insect Biochem Mol Biol 40, 405–414. doi: 10.1016/j.ibmb.2010.03.006

Perez-Hedo, M., Rivera-Perez, C. and Noriega, F. G. (2013). The insulin/TOR signal transduction pathway is involved in the nutritional regulation of juvenile hormone synthesis in *Aedes aegypti*. Insect Biochem Mol Biol 43, 495–500. doi: 10.1016/j.ibmb.2013.03.008

Raikhel A. (2005). Vitellogenesis of disease vectors, from physiology to genes. In Biology of Disease Vectors (ed. W Marquardt), pp. 329–46. London: Elsevier

Roy, S., Saha, T. T., Zou, Z. and Raikhel, A. S. (2018). Regulatory Pathways Controlling Female Insect Reproduction. Annu Rev Entomol 63, 489–511. doi: 10.1146/annurev-ento-020117-043258

Roy, S. G., Hansen, I. A. and Raikhel, A. S. (2007). Effect of insulin and 20-hydroxyecdysone in the fat body of the yellow fever mosquito, *Aedes aegypti*. Insect Biochem Mol Biol 37, 1317–1326. doi: 10.1016/j.ibmb.2007.08.004

Sheng, Z. T., Xu, J. J., Bai, H., Zhu, F. and Palli, S. R. (2011). Juvenile Hormone Regulates Vitellogenin Gene Expression through Insulin-like Peptide Signaling Pathway in the Red Flour Beetle, *Tribolium castaneum*. J Bio Chem 286, 41924–41936. doi: 10.1074/jbc.M111.269845

Shinoda, T. and Itoyama, K. (2003). Juvenile hormone acid methyltransferase: A key regulatory enzyme for insect metamorphosis. P Natl Acad Sci USA 100, 11986–11991. doi: 0.1073/pnas.2134232100

Shukla, J. N. and Nagaraju, J. (2010). Two female-specific DSX proteins are encoded by the sexspecific transcripts of dsx, and are required for female sexual differentiation in two wild silkmoth species, *Antheraea assama* and *Antheraea mylitta* (Lepidoptera, Saturniidae). Insect Biochem Mol Biol 40, 672–682. doi: 10.1016/j.ibmb.2010.06.008

Shukla, J. N. and Palli, S. R. (2012). Doublesex target genes in the red flour beetle, *Tribolium castaneum*. Sci Rep 2, doi: 10.1038/srep00948

Smykal, V., Bajgar, A., Provaznik, J., Fexova, S., Buricova, M., Takaki, K., Hodkova, M., Jindra, M. and Dolezel, D. (2014). Juvenile hormone signaling during reproduction and development of the linden bug, *Pyrrhocoris apterus*. Insect Biochem Mol Biol 45, 69–76. doi: 10.1016/j.ibmb.2013.12.003

Smykal, V. and Raikhel, A. S. (2015). Nutritional control of insect reproduction. Curr Opin Insect Sci 11, 31–38. doi: 10.1016/j.cois.2015.08.003

Song, J., Wu, Z., Wang, Z., Deng, S. and Zhou, S. (2014). Kruppel-homolog 1 mediates juvenile hormone action to promote vitellogenesis and oocyte maturation in the migratory locust. Insect Biochem Mol Biol 52, 94–101. doi: 10.1016/j.ibmb.2014.07.001

Verhulst, E. C. and van de Zande, L. (2015). Double nexus-Doublesex is the connecting element in sex determination. Brief Funct Genomics 14, 396–406. doi: 10.1093/bfgp/elv005

Wu, Q. and Brown, M. R. (2006). Signaling and function of insulin-like peptides in insects. Annu Rev Entomol 51, 1–24. doi: 10.1146/annurev.ento.51.110104.151011

Wu, Z. X., Guo, W., Xie, Y. T. and Zhou, S. T. (2016). Juvenile Hormone Activates the Transcription of Cell-division-cycle 6 (Cdc6) for Polyploidy-dependent Insect Vitellogenesis and Oogenesis. J Bio Chem 291, 5418–5427. doi: 10.1074/jbc.M115.698936

Wu, Z. X., Guo, W., Yang, L. B., He, Q. J. and Zhou, S. T. (2018). Juvenile hormone promotes locust fat body cell polyploidization and vitellogenesis by activating the transcription of Cdk6 and E2f1. Insect Biochem Mol Biol 102, 1–10. doi: 10.1016/j.ibmb.2018.09.002

